# Complementary roles for parvalbumin and somatostatin interneurons in the generation of hippocampal gamma oscillations

**DOI:** 10.1101/595546

**Authors:** Pantelis Antonoudiou, Yu Lin Tan, Georgina Kontou, A. Louise Upton, Edward O. Mann

**Affiliations:** Department of Physiology, Anatomy and Genetics, University of Oxford, Oxford, OX1 3PT, UK; Oxford Ion Channel Initiative, University of Oxford, OX1 3PT, Oxford, UK; Neuroscience, Physiology and Pharmacology, University College London

## Abstract

Gamma-frequency oscillations (30-120 Hz) can be separated into fast (>60 Hz) and slow oscillations, with different roles in neuronal encoding and information transfer. While synaptic inhibition is important for synchronization across the gamma-frequency range, the role of distinct interneuronal subtypes in fast and slow gamma states remains unclear. Here, we used optogenetics to examine the involvement of parvalbumin (PV+) and somatostatin (SST+) expressing interneurons in gamma oscillations in the mouse hippocampal CA3 *ex vivo*. Disrupting either PV+ or STT+ interneuron activity, via either photo-inhibition or photo-excitation, led to a decrease in the power of cholinergically-induced slow gamma oscillations. Furthermore, photo-excitation of SST+ interneurons induced fast gamma oscillations, which depended on both synaptic excitation and inhibition. Our findings support a critical role for both PV+ and SST+ interneurons in slow hippocampal gamma oscillations, and further suggest that STT+ interneurons are capable of switching the network between slow and fast gamma states.

## Introduction

Gamma oscillations (30 - 120 Hz) are a common feature of active cortical networks, which have been proposed to contribute to local gain control (Sohal *et al*., 2009; Cardin *et al*., 2009; Sohal, 2016) and facilitate transmission between synchronised neuronal assemblies (Fries, 2005; Akam & Kullmann, 2010; Fries, 2015). While the function of gamma oscillations remains debated (Burns, Xing & Shapley, 2011; Butler & Paulsen, 2014; Bastos, Vezoli & Fries, 2015; Ray & Maunsell, 2015; Womelsdorf & Everling, 2015; Lasztóczi & Klausberger, 2016; Sohal, 2016), changes in these rhythms continue to act as a useful marker of function and dysfunction in cortical circuit operations (Bragin *et al*., 1995; Fries *et al*., 2001; Herrmann & Demiralp, 2005; Uhlhaas & Singer, 2006; Basar-Eroglu *et al*., 2007; Uhlhaas & Singer, 2010; Yamamoto *et al*., 2014; Spellman *et al*., 2015). There is a general consensus that the generation of gamma rhythms depends upon the spiking of inhibitory interneurons, which synchronise the firing of excitatory pyramidal cells via fast synaptic inhibition (Whittington, Traub & Jefferys, 1995; Penttonen *et al*., 1998a; Csicsvari *et al*., 2003; Hajos, 2004; Mann *et al*., 2005; Hasenstaub *et al*., 2005; Bartos, Vida & Jonas, 2007; Buzsáki & Wang, 2012; Kim *et al*., 2016; Chen *et al*., 2017; Veit *et al*., 2017). Specifically, parvalbumin-expressing (PV+) interneurons, which target the perisomatic domain of pyramidal neurons, are thought to play the key role in generating and maintaining gamma oscillations in the brain (Csicsvari *et al*., 2003; Hajos, 2004; Mann *et al*., 2005; Gloveli *et al*., 2005; Hájos & Paulsen, 2009; Tukker *et al*., 2013; Cardin, 2016; Penttonen *et al*., 1998b). PV+ interneurons are adapted for fast synchronisation of network activity, as they resonate at gamma frequencies and exert strong perisomatic inhibition that is capable of precisely controlling spike timing (Pike *et al*., 2000; Pouille & Scanziani, 2001; Cardin *et al*., 2009; Bartos & Elgueta, 2012; Hu, Gan & Jonas, 2014; Kohus *et al*., 2016). Moreover, at least in the CA3 hippocampal subfield, the gamma oscillations recorded in the local field potential appear to directly reflect rhythmic perisomatic inhibitory currents (Mann *et al*., 2005; Oren, Hájos & Paulsen, 2010).

Recently, a selective role for PV+ interneurons in gamma-frequency synchronisation has been challenged by several studies performed in the primary visual cortex (Chen *et al*., 2017; Veit *et al*., 2017; Hakim, Shamardani & Adesnik, 2018). In this brain region, it was shown that dendrite-targeting somatostatin-expressing (SST+) interneurons were the main contributors for the generation of slow gamma oscillations, while PV+ interneurons were more important for higher frequency synchronisation (Chen *et al*., 2017). Previous studies have found analogous roles for SST+ and PV+ interneurons in low- and high-frequency network synchronisation (Beierlein, Gibson & Connors, 2000; Gloveli *et al*., 2005; Tukker *et al*., 2007; Craig & McBain, 2015). However, it is not yet clear if it is the frequency tuning of each interneuronal circuit that varies across brain areas, or whether SST+ interneurons might play a more generic role in the generation of slow gamma oscillations.

The hippocampus displays both slow and fast gamma rhythms during theta activity, with slow gamma generated in CA3 and fast gamma propagated from entorhinal cortex (Bragin *et al*., 1995; Colgin *et al*., 2009; Schomburg *et al*., 2014; Lasztóczi & Klausberger, 2016). The circuitry for slow gamma oscillations is preserved in hippocampal slices (Fisahn *et al*., 1998), and these models have been used extensively to show that PV+ interneurons are strongly phase-coupled to gamma oscillations, and contribute to rhythmogenesis (Hajos, 2004; Mann *et al*., 2005; Gloveli *et al*., 2005; Gulyás *et al*., 2010). However, the majority of interneurons are phase-coupled to ongoing slow gamma oscillations (Hajos, 2004; Gloveli *et al*., 2005; Oren *et al*., 2006), and it may be that SST+ interneurons play an important role in synchronising PV+ networks. Indeed, whether specific classes of CA3 interneuron are necessary and sufficient for the generation of slow gamma oscillations has not yet been tested. Here, we took advantage of optogenetic techniques (Nagel *et al*., 2003; Chow *et al*., 2010; Boyden *et al*., 2005) to test the involvement of PV+ and SST+ interneurons in cholinergically-induced gamma oscillations in the CA3 of acute hippocampal slices.

## Results

### PV+ interneuron activity is necessary for cholinergically-induced gamma oscillations in hippocampal CA3

In order to test if the activity of PV+ interneurons is necessary for the generation of slow hippocampal gamma oscillations, we took advantage of optogenetic photo-inhibition (Chow *et al*., 2010). We injected PV-cre mice with AAV carrying the inhibitory proton pump archaerhodopsin (Arch3-eYFP or ArchT-GFP). Expression of Arch in PV-cre mice was restricted to the pyramidal cell layer indicating selective expression in perisomatic targeting PV+ interneurons (Fig. 1a) (Somogyi & Klausberger, 2005; Royer *et al*., 2012; Hu, Gan & Jonas, 2014). Intracellular recordings performed in opsin expressing cells demonstrated that these cells were fast-spiking and that sustained light illumination was able to produce robust hyperpolarisation, indicating functional expression of Arch in PV+ interneurons (Supplementary Fig. 1d-e).

**Figure 1:**
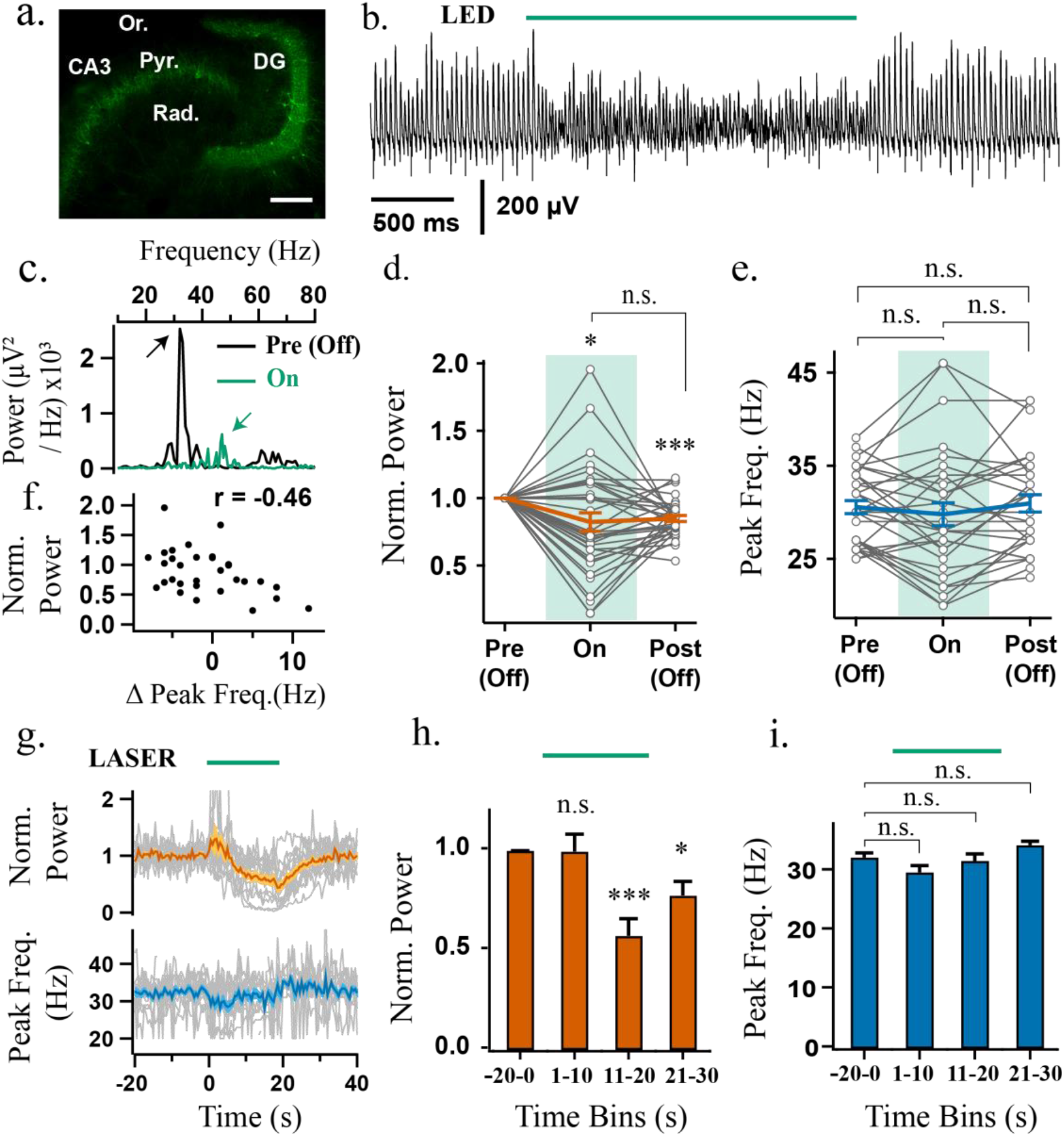
Sustained photo-inhibition of PV+ interneurons suppresses the power of gamma oscillations. a-e) photo-inhibition with LED and g-i) photo-inhibition with laser experiments. a) Confocal image of ventral hippocampus slice from a PV-cre mouse injected intrahippocampally with AAV-Arch3 eYFP. CA3 = Cornu Ammonis 3, DG = Dentate Gyrus, Pyr. = stratum Pyramidale, Rad. = stratum Radiatum, Or. = Stratum Oriens. Scale bar = 200 μm. b) LFP recording from CA3 stratum pyramidale illustrating effect of PV+ interneuron photo-inhibition (LED, 530 nm, approx. 4.25 mW) on gamma oscillations induced by the application of 5 μM Cch. c) Representative power spectra before (black) and during (green) LED illumination (arrows indicate power spectrum peaks). d) Power area in the 20-100 Hz band normalised to baseline (Pre (Off)) during (On) and after LED stimulation (Off (Post)) (n = 35). e) Peak frequency for experiments when the oscillation was not abolished (n = 31/35). f) Power area change versus peak frequency difference recorded between stimulation and baseline periods. g) Stronger photoinhibition was achieved using high power laser illumination (approx. 18.6 mW). Top: Change in power area normalised to baseline. Bottom: Peak frequency of the oscillation calculated in 1 second bins across experiments (n = 14). h) Mean change in power area normalised to baseline (n = 14). i) Mean peak frequency for trials when the oscillation was not abolished (n = 13). *p < 0.05, **p < 0.01, ***p < 0.001, n.s. p >= 0.05. Changes in peak frequency were analysed using rmANOVA, followed by post-hoc paired t-tests with correction for multiple comparisons. Solid brackets represent paired t-tests and standalone star symbols represent one-sample t-test versus normalised baseline. Grey lines represent single experiments. Error bars and shaded area are SEM and coloured line the population mean.

Gamma oscillations were induced in hippocampal slices from PV-Arch mice in area CA3 using bath application of the cholinergic agonist carbachol (Cch - 5 μM). Local field potential recordings from the CA3 pyramidal cell layer revealed robust gamma oscillations that were centred around 30 – 40 Hz with clear side peaks in the autocorrelogram (Supplementary Fig. 1a-c), as has been reported previously (Fisahn *et al*., 1998; Hajos, 2004; Mann *et al*., 2005). Overall, sustained photo-inhibition of PV+ interneurons using LED illumination (< 5mW) significantly decreased gamma power area (0.82 +/- 0.068 of baseline period, t = 2.59, p = 0.029, one sample t-test; Fig. 1 b-d), although increases in power were observed in some slices (Fig. 1d). A significant suppression was also observed in the period of 0.5 - 1.5 seconds following light illumination termination (0.85 +/- 0.022 of baseline period, t = 6.70, p < 0.001, one sample t-test; Fig. 1d). However, the light-induced changes in gamma power were reversible, as there were no significant changes in the gamma power area recorded during the baseline periods across trials (F(4, 116) = 0.68, p = 0.61, rmANOVA). The changes in gamma power were not accompanied by a consistent change in gamma frequency (F(1.44, 43.23) = 1.25, p = 0.288, rmANOVA; Fig. 1e), although there was a significant correlation between the changes in frequency and power area (t = 2.77, p = 0.01, Pearson correlation, Fig. 1f), suggesting a consistent disturbance to endogenous oscillatory activity.

While LED photo-inhibition of PV+ interneurons significantly modulated gamma power, the oscillations did not collapse. Pyramidal neurons make strong recurrent connections with PV+ interneurons (Mann, Radcliffe & Paulsen, 2005; Oren *et al*., 2006; Hofer *et al*., 2011; Packer & Yuste, 2011; Bartos & Elgueta, 2012; Kohus *et al*., 2016), and it might be hard to break these feedback loops with photo-induced inhibitory currents. To test this possibility, we used long-lasting laser illumination with the prospect of biochemically silencing PV+ interneurons, by preventing synaptic release via terminal alkalisation (El-Gaby *et al*., 2016). PV+ interneurons expressing ArchT-GFP were illuminated with sustained green laser light (532 nm, approx. 18 mW for 20 seconds). Similar to the LED experiments, there were inconsistent network responses to PV+ interneuron photo-inhibition at the beginning of laser illumination (1.00 +/- 0.087 of baseline period, t = 0.04, p = 1.00, one sample t-test; Fig. 1g&h). However, the power of the oscillation consistently decreased during sustained laser illumination (0.57 +/- 0.086 of baseline period, t = 5.00, p < 0.001, one sample t-test; Fig. 1g&h) and remained suppressed in the light-off period following laser stimulation (0.78 +/- 0.071 of baseline period, t = 3.17, p = 0.022, one sample t-test; Fig. 1g, h). There was no consistent effect on the frequency of the oscillations (F(1.92, 23.08) = 7.77, p = 0.003, rmANOVA; Fig. 1g&i). Laser illumination of PV+ interneurons expressing only control fluorophore did not alter gamma oscillation power nor frequency (Supplementary Fig. 1g-i). This slow and selective process of decreasing gamma power is consistent with biochemical silencing of synaptic terminals (El-Gaby *et al*., 2016). These results further support the importance of PV+ interneuron activity in generating gamma oscillations in hippocampal area CA3 (Hajos, 2004; Mann *et al*., 2005; Gulyás *et al*., 2010; Tukker *et al*., 2013). Residual gamma oscillations following photo-inhibition of PV+ interneurons may reflect incomplete transfection of the PV+ network or the presence of a distinct oscillatory circuit.

### SST+ interneurons are necessary for Cch-induced gamma oscillations in hippocampal area ca3

To examine if SST+ interneuron activity is also required during Cch-induced gamma oscillations in CA3, we injected the AAV-Arch vector (Arch3-eYFP or ArchT-GFP) intrahippocampally in SST-cre mice. Expression of Arch was restricted to the strata oriens, radiatum and lacunosum moleculare (Fig. 2a), suggesting expression in SST+ dendrite-targeting interneurons (Ma *et al*., 2006; Lovett-Barron *et al*., 2012; Muller & Remy, 2014; Urban-Ciecko & Barth, 2016). Whole-cell recordings were performed in opsin positive cells and indicated functional expression of Arch (n = 4, Supplementary Fig. 2a-b).

**Figure 2:**
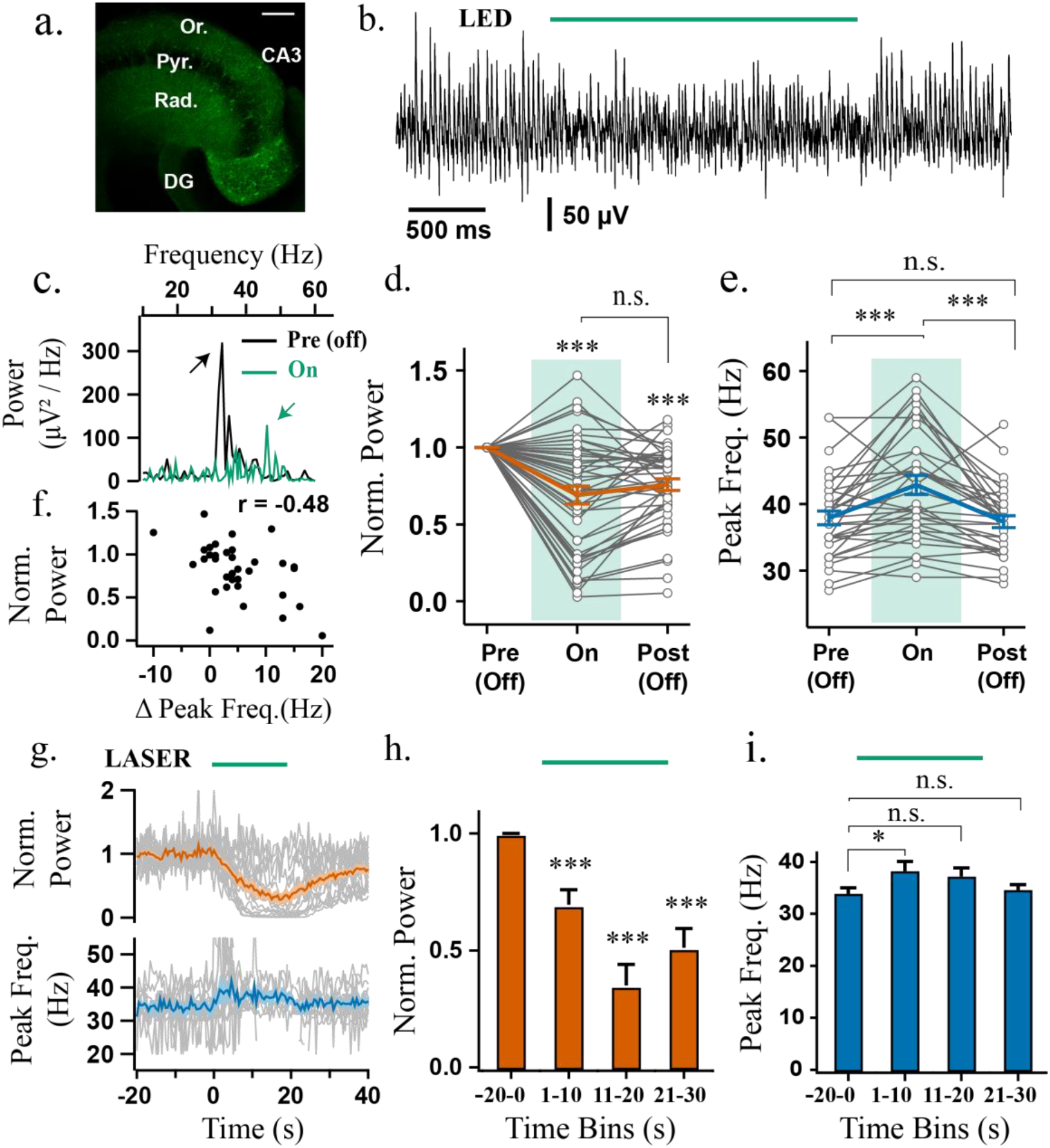
Sustained photo-inhibition of SST+ interneurons suppresses gamma power and increases frequency. a) Confocal image of ventral hippocampus slice from SST-cre mice with eYFP-Arch3 expression. CA3 = Cornu Ammonis 3, DG = Dentate Gyrus, Pyr. = stratum Pyramidale, Rad. = stratum Radiatum, Or. = Stratum Oriens. Scale bar = 200 μm. b) Representative LFP recordings from CA3 area illustrating effect of SST+ interneuron photo-inhibition (LED, 530 nm, approx. 4.25 mW) on gamma oscillations along with c) its respective power spectrum (arrows indicate power spectrum peaks). d) Power area in the 20 −100 Hz band normalised to baseline (Pre (Off)) during (On) and after LED stimulation (Post (Off)) (n = 44). e) Peak frequency for experiments when the oscillation was not abolished (n = 33/44). f) Power area change against peak frequency difference between stimulation and baseline periods. g) Top: Change in power-area normalised to baseline and Bottom: Peak frequency of the oscillation calculated in 1 second bins across experiments with high-power laser stimulation (approx. 18.6 mW; n = 17). h) Average power change normalised to baseline (n = 17). i) Average peak frequency (n = 11/17). *p < 0.05, **p < 0.01, ***p < 0.001, n.s. p >= 0.05. Changes in peak frequency were analysed using rmANOVA, followed by post-hoc paired t-tests with correction for multiple comparisons. Solid brackets represent paired t-tests and standalone star symbols represent one-sample t-test versus normalised baseline. Grey lines represent single experiments, error bars and shaded area are SEM and coloured line the population average.

Unlike the experiments with PV+ photo-inhibition, sustained photo-inhibition of SST+ interneurons using LED illumination (< 5mW) reliably decreased gamma oscillation power (0.69 +/- 0.057 of baseline period, t = 5.40, p < 0.001, one sample t-test; Fig. 2c, d), which remained suppressed in the immediate period following SST+ interneuron photoinhibition (0.76 +/- 0.039 of baseline period, t = −6.26745, p < 0.001, one sample t-test; Fig. 1d). This post-light suppression was reversed from trial to trial (F(4, 164) = 2046, p = 0.048, rmANOVA; all paired t-tests t > 2.81, p > 0.07). In addition, light stimulation significantly modulated oscillation frequency (F(1.25, 39.97) = 22.60, p < 0.001, rmANOVA), with an increase in frequency from 37.79 +/- 1.083 Hz to 43.00 +/- 1.466 Hz during light stimulation (t = 4.74, p < 0.01, paired t-test), which reversed following light offset (Fig. 2e).

Stronger laser illumination in the first half of the stimulation period (532 nm, approx. 18 mW for 20 s) had similar effects as the LED experiment. Specifically, the power of Cch gamma oscillations decreased (0.70 +/- 0.064 of baseline, t = 4.76, p < 0.01, one-sample t-test), and the peak frequency increased (34.22 +/- 1.191 Hz to 38.60 +/- 1.868 Hz, t = 3.93, p = 0.017, paired t-test; rmANOVA, F(1.36, 13.63) = 5.47, p = 0.027; Fig. 2g-i). During the second half of the stimulation period, gamma power was strongly suppressed (0.35 +/- 0.090 of baseline, t = 7.23, p < 0.01, one sample t-test; Fig. 2g-h), often resulting in oscillation collapse (7/13 slices). This could indicate that silencing SST+ interneurons is sufficient to disrupt the hippocampal network during gamma oscillations and that SST+ interneuron activity is necessary for proper maintenance of Cch-induced oscillations in the CA3 area of the hippocampus. Moreover, the frequency of the Cch-gamma oscillations remains upregulated for the whole duration of laser illumination when the oscillations do not collapse (Fig. 2i vs Supplementary Fig.2d), but in each case remained below 60 Hz, suggesting that SST+ interneurons can exert strong control over the frequency of slow gamma oscillations.

### Rhythmic synchronisation of the hippocampal network by perisomatic and dendritic inhibition

The experiments using photo-inhibition indicate that the generation of gamma oscillations in hippocampal area CA3 involves the endogenous recruitment of both PV+ and SST+ interneurons. In order to test whether the activation of PV+ or SST+ interneurons is sufficient to synchronise the hippocampal network at gamma frequencies, we next examined cell type-specific photo-stimulation using Channelrhodopsin 2 (ChR2) (Nagel *et al*., 2003; Boyden *et al*., 2005). Injection of AAV-ChR2-mCherry produced similar expression patterns as Arch in both PV- and SST-Cre mouse lines (Fig. 3a, b). Photo-stimulation of ChR2-expressing PV+ interneurons at 40 Hz (1 ms pulse width) entrained ongoing oscillations in 14/18 experiments (>2mW; n = 12 at 5.5 mW, n = 6 at 2.2 mW - merged due to similar effects; Fig. 3c, f and Supplementary Fig. 3ai). In the remaining 4 out of 18 experiments the ongoing oscillations were not entrained (Fig. 3f; Supplementary Fig. 3aiii). This effect was likely observed due to low ChR2 expression, as pulses with longer width (5 ms) entrained the oscillation in the same experiments (Supplementary Fig. 3b-c). Thus, PV+ interneurons are sufficient to synchronise the hippocampal network at gamma frequencies.

**Figure 3:**
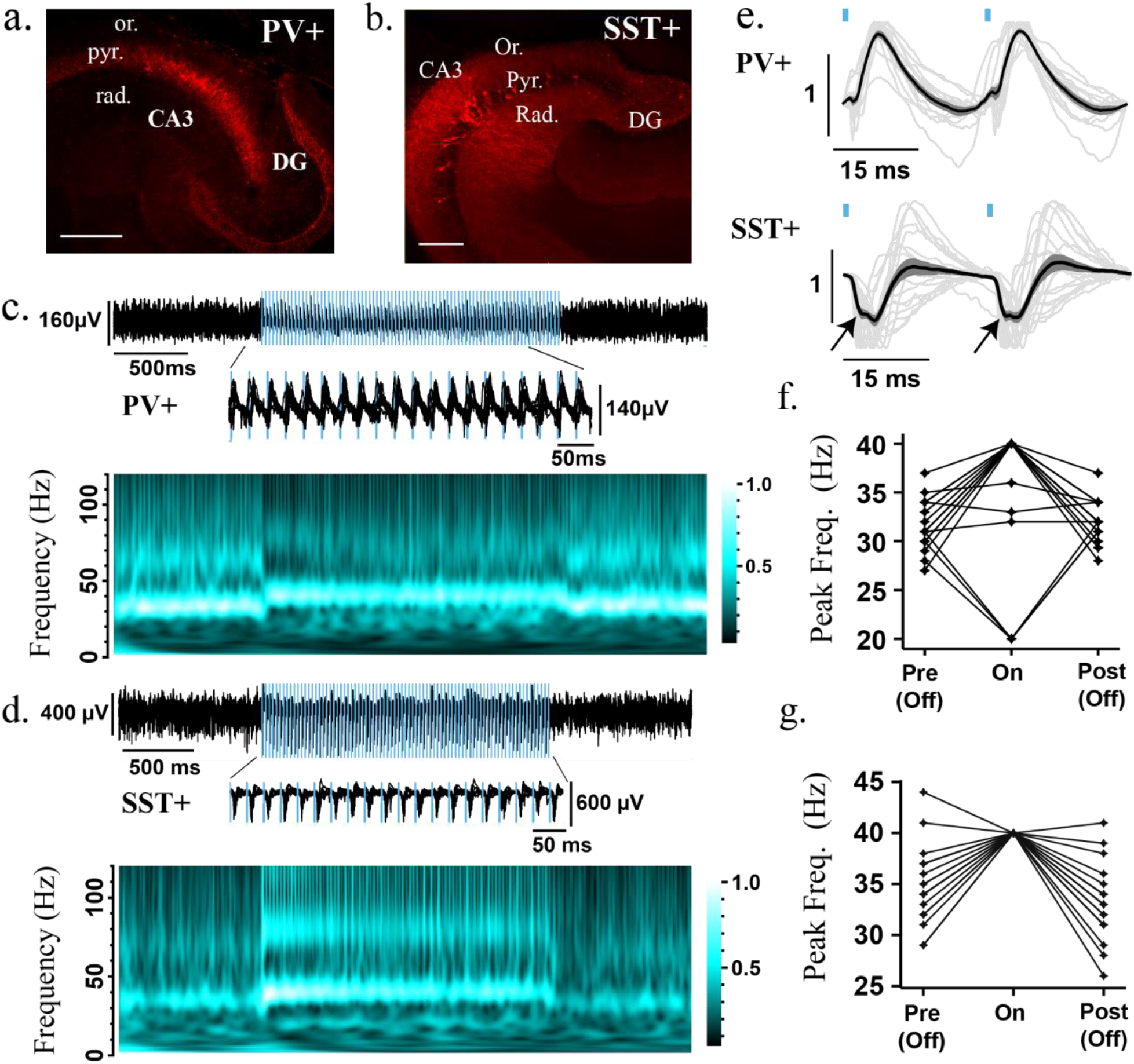
Rhythmic photo-stimulation of either PV+ or SST+ interneurons entrains Cch-induced gamma oscillations. Confocal image of ventral hippocampus (350 μm slice) from a: PV-cre mice and b: SST-cre mice with mCherry-ChR2 expression. CA3 = Cornu Ammonis 3, DG = Dentate Gyrus, Pyr. = stratum Pyramidale, Rad. = stratum Radiatum, Or. = Stratum Oriens. Scale bar = 200 μm. c and d) Top: Representative LFP recording from CA3 area illustrating the entrainment of Cch-induced oscillations to 40 Hz light pulses in c: PV-cre and d: SST-cre mice expressing mCherry-ChR2 (1ms pulse width; blue light illumination at 5.5 mW). Middle: Zoomed recordings during pulse stimulation. Bottom: Magnitude component of the wavelet transform normalised by its maximum value. Brighter colours represent larger gamma power. e) Normalised average waveform following two consecutive pulses at 40 Hz and 1 ms pulse width from each experiment (PV+: n = 15/18; SST+: n = 19/22). Black line is the population average, grey lines represent individual experiments and dark-grey shaded area is the SEM Black. Arrows indicate initial negative peak. f, g) Peak frequency of oscillation before (Pre (Off)), during (On) and after (Post (Off)) light stimulation in f: PV+ (n = 18/18) and g: SST+ (n = 19/22) experiments (note experiments entrained at 20 Hz reflect suppression of alternate gamma cycle – see suppl. Fig. 3aii). Black lines represent single experiments.

Rhythmic photo-stimulation of SST+ interneurons reliably entrained ongoing oscillations in 19 out of 22 experiments (>2mW; n = 13 at 5.5 mW, n = 9 at 2.2 mW - merged due to similar effects; Fig. 3d, g). In the remaining 3 out of 22 oscillations were abolished during 40 Hz photo-stimulation. These results indicate that transient activation of SST+ dendrite-targeting interneurons is also sufficient to synchronise the hippocampal network at gamma frequencies. Activation of PV+ and SST+ interneurons produced opposite deflections in the pulse-locked waveform of the LFP recorded in the stratum pyramidale (Fig. 3c-e), as might be expected from the somatodendritic profile of their axon terminations. However, activation of SST+ interneurons was sometimes accompanied by an initial fast negative component (Fig. 3e), which was reminiscent of a population spike arising from the synchronised firing of excitatory cells in the hippocampus (Andersen, Bliss & Skrede, 1971; Wierenga & Wadman, 2003), despite the sparsity of SST+ axons in this layer.

To study the SST+ induced waveform in isolation, we repeated the same experiment in quiescent slices, perfused only with aCSF. Blue light pulses (1 ms width) at 40 Hz induced strong pulse-locked field responses with fast-negative deflections, which were resistant to glutamate receptor blockers (Supplementary Fig. 3d-f), but were followed by a glutamate receptor-mediated positive deflection. The fast-negative deflections did not appear to reflect fast GABAergic transmission, as application of GABA_A_ receptor (GABA_A_R) blockers lead to light-induced epileptiform bursts (n = 4; Supplementary Fig. 3g). These results suggest that SST+ interneuron photo-activation generates network excitation, that is not mediated through GABA_A_Rs, at the onset of light illumination. We did not observe ChR2 expression in CA3 pyramidal neurons during intracellular recordings (n = 18, supplementary Fig. 5e), although there have been reports of off-target expression in SST+ interneurons of juvenile animals (Taniguchi *et al*., 2011). An alternative possibility is that robust activation of a dense plexus of SST+ axons in the dendritic layers is sufficient to induce spiking in pyramidal neurons via ephaptic coupling (Ferenczi *et al*., 2016a). Either scenario makes it difficult to interpret the results of pulsed stimulation in the SST-ChR2 mice, but any electrically-mediated bystander effects are likely to occur during stimulus onset (maximal hypersynchrony), and may be less relevant during more sustained patterns of stimulation.

### Sustained activation of PV+ interneurons suppresses Cch-induced gamma oscillations

We used two patterns of sustained activation, light steps and fully-modulation sine waves at 8 Hz, and tested these in slices from PV-ChR2 mice. In a subset of light step experiments, we recorded ongoing gamma oscillations in the LFP whilst tonically driving PV+ interneurons at increasing strengths across trials (by changing the levels of blue light illumination, 10 - 5500 μW). The change in power between baseline and light activation period was measured at each light intensity level. We then obtained the response level at which the power changed by half of the maximum for each experiment (half-maximal response). For half-maximal response trials, photo-activation of PV+ interneurons (2 seconds) consistently decreased the power-area (0.52 +/- 0.016 compared to baseline, t = −29.56, p < 0.01, one-sample t-test; Fig. 4a-c, e) and increased the peak frequency (from 32.70 +/- 0.793 Hz (baseline) to 38.76 +/- 1.094 Hz, t = 8.21, p < 0.01, paired t-test; Fig. 4d, f). Furthermore, there was a progressive decrease in power (r = −0.84, n = 121 values, t = 17.00, p < 0.01) and increase in frequency (r = 0.49, n = 100/121values, t = 5.60, p < 0.01) as the light intensity increased (Fig. 4g-h). In order to estimate the maximal effect of PV+ interneuron stimulation, we pooled experiments using strong light intensity illumination (> 2mW, including cases where light intensity-response curves were not assessed; n = 14 at 5.5 mW, n = 9 at 2.2 mW). Overall, strong light illumination caused a substantial decrease in the normalised power-area (0.09 +/- 0.029, t = p < 0.01, one-sample t-test; Fig. 4i, j) and abolished the oscillations in most experiments (17/23). These results indicate that progressive up-regulation of PV+ interneuron activity decreases gamma power and increases the frequency until the rest of the hippocampal network is fully silenced.

**Figure 4:**
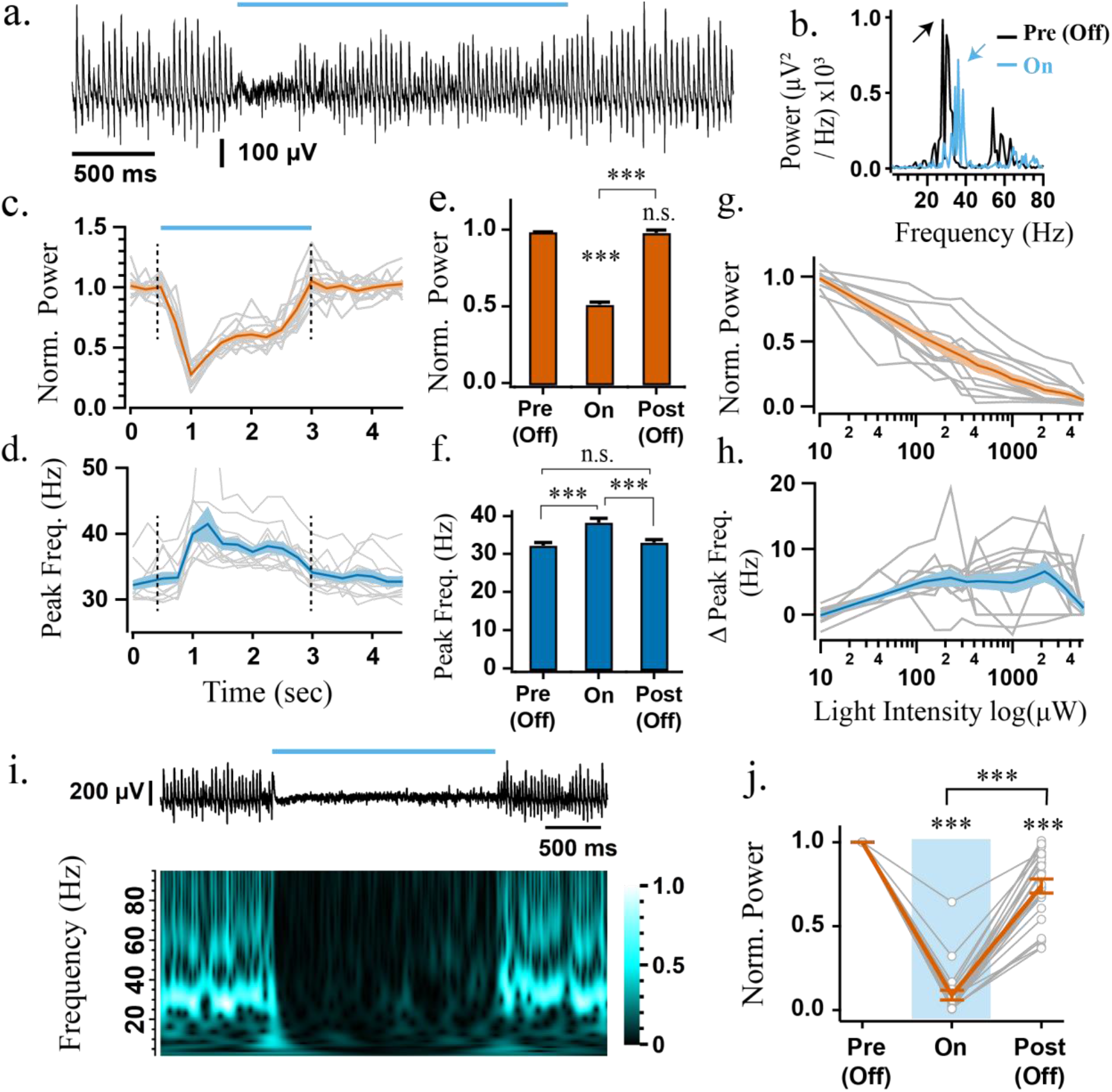
Sustained photo-excitation of PV+ interneurons decreases the power and increases the frequency of Cch-induced gamma oscillations. a) Representative LFP recordings from CA3 area illustrating effect of PV+ interneurons photo-excitation (155 μW) on gamma oscillations along with b) its respective power spectrum (arrows indicate power spectrum peaks). c) Power-area normalised to baseline and d) peak frequency of the oscillation calculated in 0.5 second bins across experiments (n = 12). e) Average change in power-area during (On) and after light stimulation (Post (Off)) normalised to baseline (Pre (Off)). f) Average peak frequency; rmANOVA: F(1.14, 12.50) = 44.14, p < 0.001. g) Power-area change and h) frequency difference between light stimulation period and baseline plotted against light intensity (n = 12). i) Top: Representative LFP recording from CA3 area illustrating the collapse of Cch-induced oscillations in response to strong and sustained blue light illumination (5.5 mW). Bottom: Magnitude component of the wavelet transform normalised by its maximum value. j) Average change in power-area upon strong and sustained blue light illumination (n total = 23 >2mW: n = 14 at 5.5 mW and n = 9 at 2.2 mW). Changes in peak frequency were analysed using rmANOVA, followed by post-hoc paired t-tests with correction for multiple comparisons. *p < 0.05, **p < 0.01, ***p < 0.001, n.s. p >= 0.05. Solid brackets represent paired t-tests and standalone star symbols represent one-sample t-test versus normalised baseline. Grey lines represent single experiments, error bars and shaded area are SEM and coloured line the population average.

Interneurons have been shown to be particularly susceptible to depolarisation block (Herman *et al*., 2014). In order to ensure that these effects were not caused from impaired action potential generation in PV+ interneurons (i.e. depolarisation block), as we recorded spiking activity using a linear multi-electrode array (MEA) during Cch-induced oscillations. PV+ interneurons (spike width: 0.49 +/- 0.04 ms) maintained spiking activity during sustained illumination (5.5 mW; Median sustained activation index [IQR] = 0.87 [0.46, 1], Z=171, p<0.001, n=18, one-sample Wilcoxon signed rank test; analysis performed on last second of trial), and this was associated with decreased activity of regular spiking (RS; −0.72 [−0.92, −0.40]; Z=2, p<0.001, n=53, one-sample Wilcoxon signed rank test; z=65.7, p<0.001 cf. PV+ interneurons, Kruskal-Wallis Test followed by posthoc Dunn’s test with Bonferroni correction for multiple comparisons) and fast-spiking cells (FS; −0.52 [−0.82, −0.12]; Z=141 p<0.001, n=49, one-sample Wilcoxon signed rank test; z=50.1, p<0.001 cf. PV+ interneurons, Kruskal-Wallis Test followed by posthoc Dunn’s test with Bonferroni correction for multiple comparisons) (Supplementary Fig.4). These results are consistent with increased PV+ interneuron activity during light illumination that leads to reduced activity in hippocampal principal cells.

During 8 Hz sinusoidal modulation of PV+ interneurons, the instantaneous gamma magnitude, assessed using the Hilbert transform, was found to be negatively correlated with light intensity in agreement to light step experiments (across all experiments, Pearson correlation, mean r = −0.51 +/- 0.04, t > 23.2, p < 0.01, n = 12) (Suppl. Fig.4d). During MEA recordings, spike rates of PV+ interneurons correlated positively with theta-frequency changes in light intensity (Median rank correlation coefficient [IQR] = 0.75 [0.55, 0.83], Z=120, p=0.001, n=15, one-sample Wilcoxon signed rank test), while negative correlations were found for the spike rates of RS (−0.19 [−0.37, −0.09]; Z=100, p<0.001, n=43, one-sample Wilcoxon signed rank test; z=48.3, p<0.001 cf. PV+ interneurons, Kruskal-Wallis Test followed by posthoc Dunn’s test with Bonferroni correction for multiple comparisons) and FS cells (−0.19 [−0.39, −0.01]; Z=232, p=0.006, n=42, one-sample Wilcoxon signed rank test; z=45.7, p<0.001 cf. PV+ interneurons, Kruskal-Wallis Test followed by posthoc Dunn’s test with Bonferroni correction for multiple comparisons). These findings indicate that a transient increase in PV+ interneuron activity causes a rapid and reversible decrease in the power of the Cch-gamma oscillations and firing rates of other neurons.

### Sustained activation of SST+ interneurons induces fast gamma oscillations

We obtained the light intensity response curves with light steps in slices from SST-ChR2 mice and observed similar results as in PV-ChR2 experiments. Sustained light illumination decreased the power (0.49 +/- 0.029, t = 17.53, p < 0.01, one sample t-test; Fig. 5a-c, e) and increased the frequency at half-maximal response (from 34.08 +/- 0.954 Hz during baseline period to 38.17 +/- 1.400 Hz, t = 3.658, p = 0.011, paired t-test; Fig. 5d, f). Moreover, as the light intensity increased, the power progressively decreased (r = −0.66, n = 107 values, t = 9.11, p < 0.01; Fig. 5g), and frequency progressively increased (r = 0.71, n = 56 out of 107 values, t = 7.41, p < 0.01; Fig. 5h). It is perhaps not surprising that excitatory networks can be suppressed by photo-activation of GABAergic interneurons. However, different responses were revealed when we assessed the effects of strong photo-activation of SST+ interneurons on Cch-induced gamma oscillations (light-intensity response curves where performed in a subset of experiments; n = 18 slices at 5.5 mW and n = 13 slices at 2.2 mW, merged). Consistent with PV-ChR2 step experiments, the gamma power was reduced during light stimulation when compared to baseline period (0.34 +/- 0.150, t = −4.39, p < 0.01, one sample t-test; Fig. 4.6b) and in approximately half of the experiments, gamma oscillations were fully abolished (n = 16/31 slices). In contrast, in experiments where the oscillations persisted, their frequency increased strongly from 34.63 +/- 0.836 Hz during baseline to 62.75 +/- 4.921 Hz during light illumination (n = 15/31 slices; t =5.61, p < 0.01, paired t-test; Fig 5i-k). These fast gamma oscillations occurred most reliably in slices that the light-intensity response curves were not obtained. In order to test if SST+ interneuron photo-activation alone is sufficient to induce oscillations, as opposed to simply increasing the frequency of ongoing activity, we repeated the same experiments in the absence of Cch. Sustained photo-activation of SST+ interneurons induced *de novo* oscillations in the fast gamma-band range with peak frequency of 80.5 +/- 2.48 Hz (12/16 slices; Fig. 6a, c). Isolating the CA3 area from DG did not prevent the generation of *de novo* oscillations (n = 3 slices).

**Figure 5:**
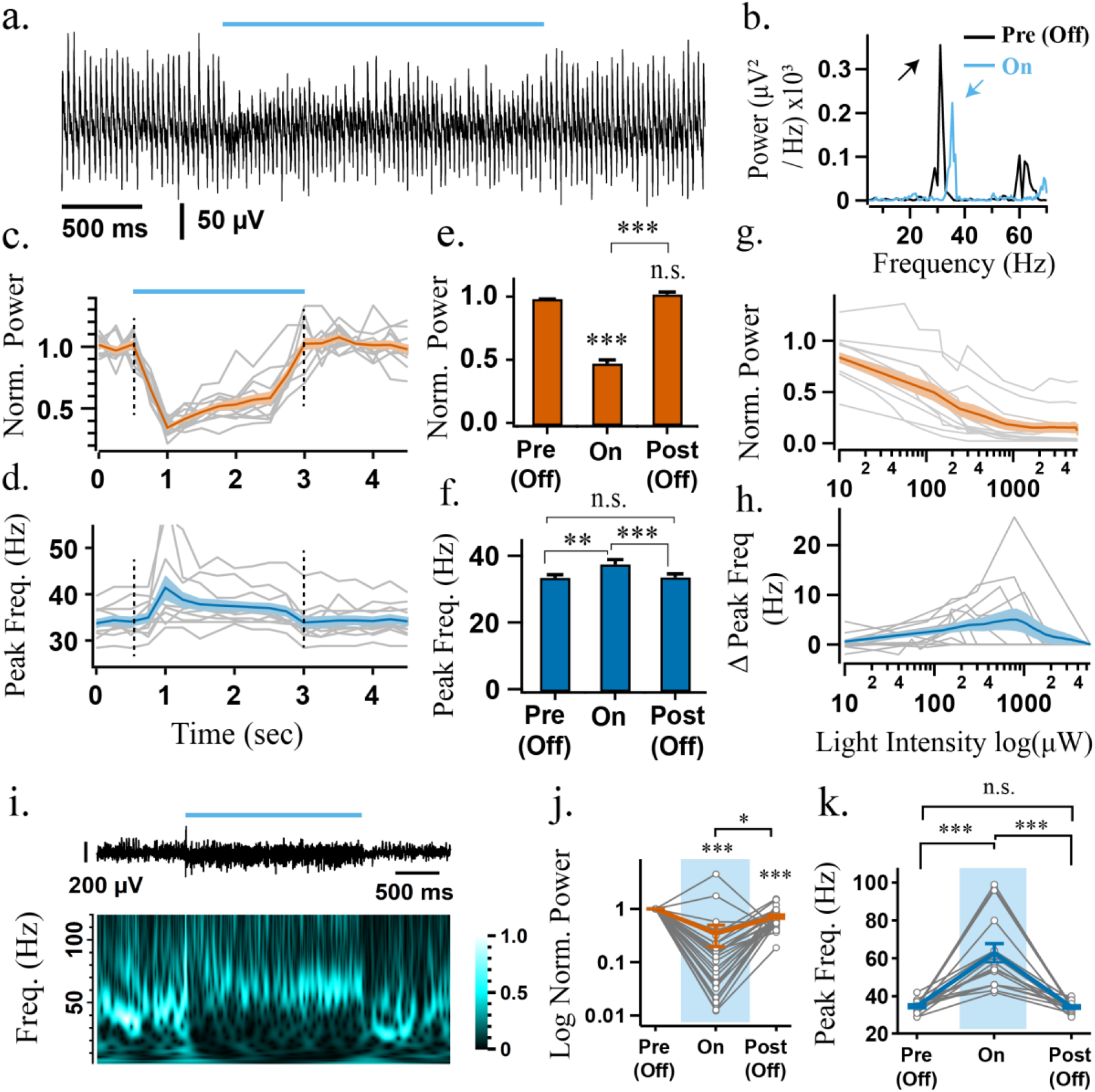
Sustained photo-excitation of SST+ interneurons decreases the power and increases the frequency of Cch-induced gamma oscillations but can also induce high-frequency oscillations. a) Representative LFP recordings from CA3 area illustrating effect of SST+ interneurons photo-excitation (155 μW) on gamma oscillations along with b) its respective power spectrum (arrows indicate power spectrum peaks). c) Power-area normalised to baseline and d) peak frequency of the oscillation calculated in 0.5 second bins across experiments (n = 12). e) Average change in power-area during (On) and after light stimulation (Post (Off)) normalised to baseline (Pre (Off)). f) Average peak frequency; rmANOVA: F(1.05, 11.59) = 15.05, p = 0.002. g) Power-area change and h) frequency difference between light stimulation period and baseline plotted against light intensity (n = 12). i) Top: Representative LFP recording from CA3 area illustrating the induction of high-frequency oscillations in response to strong and sustained blue light illumination (5.5 mW). Bottom: Magnitude component of the wavelet transform normalised by its maximum value. j) Normalised power of Cch-oscillations during SST+ interneurons cell photo-activation (n = 31). k) Peak frequency of oscillations that were not abolished from strong light illumination (n remaining = 16/31: n = 4 at 5.5 mW and n = 12 at 2.2 mW); rmANOVA: F(1.03, 15.39) = 31.45, p < 0.001.. Changes in peak frequency were analysed using rmANOVA, followed by post-hoc paired t-tests with correction for multiple comparisons. *p < 0.05, **p < 0.01, ***p < 0.001, n.s. p >= 0.05. Solid brackets represent paired t-tests and standalone star symbols represent one-sample t-test versus normalised baseline. Grey lines represent single experiments, error bars and shaded area are SEM and coloured line the population average.

**Figure 6:**
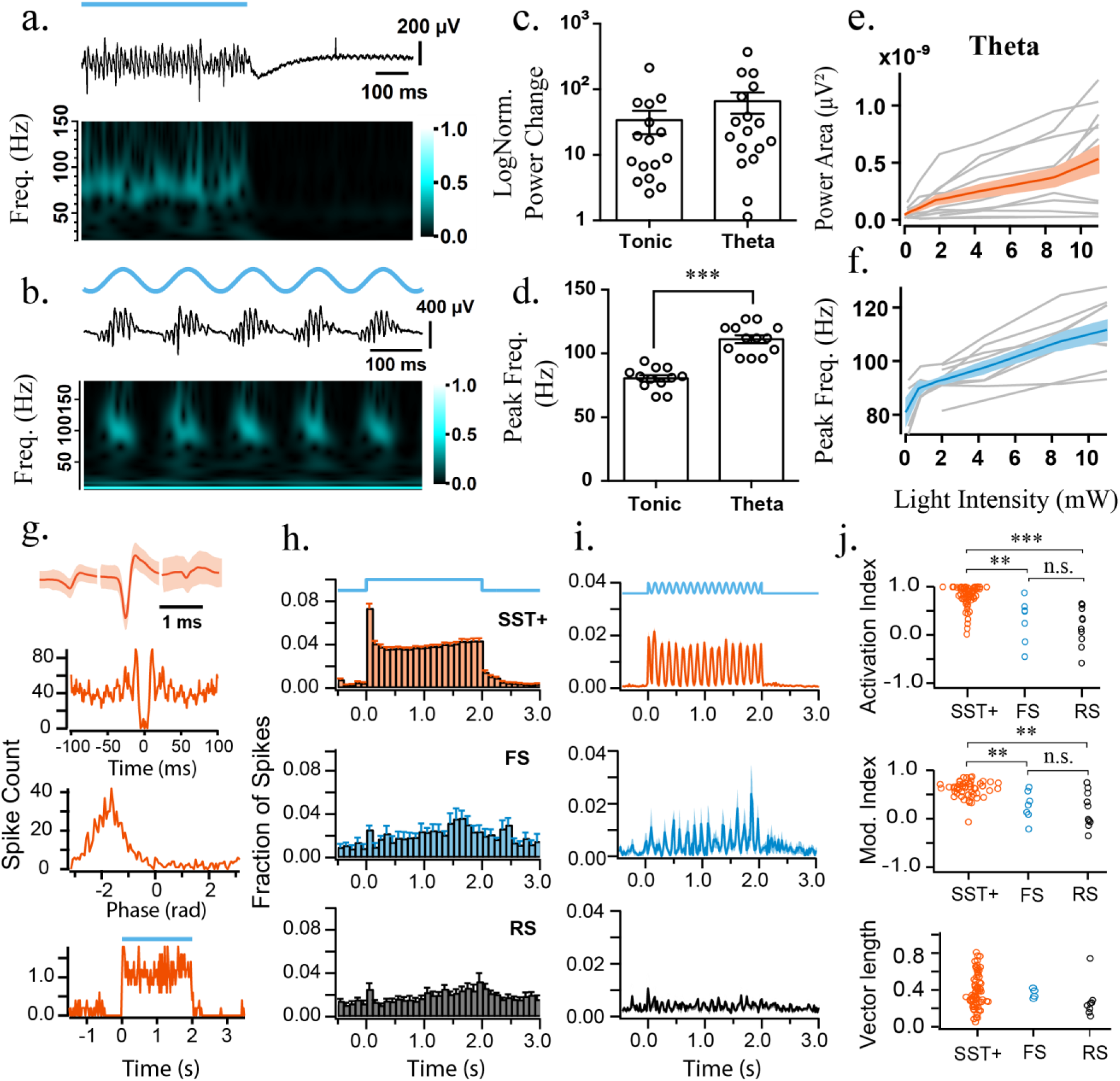
Photo-activation of SST+ interneurons induces *de novo* oscillations in the absence of Cch. a, b) Top: Representative LFP recording from CA3 area illustrating induction of high-frequency oscillations by a) constant and b) theta blue light illumination (10 mW). Bottom: Magnitude component of the wavelet transform normalised by its maximum value; brighter colours represent larger magnitude. c) Log power change compared to baseline during constant (n = 16) and theta blue light illumination (n = 17). d) Peak frequency of the *de novo* oscillations is higher when induced by theta when compared to tonic stimulation; two-sample t-test, ***p < 0.001. Grey lines/markers represent single experiments, black line is the population average and error bars are the SEM. e) Power-area and f) frequency during light stimulation period plotted against light intensity of theta photo-activation (n = 12). The black line is the average response and the dark-grey shaded area represents SEM. (g-i) Multi-unit recordings during SST+ interneuron sustained photo-excitation g) Putative opto-tagged SST+ average spike waveform, autocorrelation, phase and step histogram during sustained light illumination. h) step histogram, i) sinusoidal histogram of photo-tagged SST+, FS and RS cells. Shaded regions represent standard deviation. j) top to bottom: sustained activation index, modulation index and vector length. Kruskal-Wallis Test was followed by posthoc Dunn’s test with Bonferroni correction for multiple comparisons. *p < 0.05, **p < 0.01, ***p < 0.001, n.s. p >= 0.05.

Furthermore, sinusoidal light activation at 8 Hz (theta photo-activation) also induced robust oscillations with higher peak frequency than the tonic activation 111.2 +/- 3.15 Hz (13/17 slices; t = 7.64, p < 0.001, two-sample t-test; Fig. 6b, d-e). This is consistent with previous experiments showing that transient light activation induces higher frequency oscillations than sustained illumination (Butler *et al*., 2016; Betterton *et al*., 2017). Furthermore, the power (r = 0.67, n = 70 values, t = 7.52 p < 0.01) and frequency (r = 0.77, n = 48/70 values, t = 8.20, p < 0.01) of the *de novo* oscillations progressively increased as the light intensity of theta photo-activation was elevated (Fig. 6f). This monotonic increase in peak frequency contrasts with the properties of oscillations induced by photo-activation of principal cells in the hippocampus, where the frequency of the oscillations remains relatively constant within the slow gamma band across light intensities (Butler *et al*., 2016; Betterton *et al*., 2017; Butler, Hay & Paulsen, 2018). Therefore, SST+ interneuron photo-activation in CA3 appears to induce a distinct type of gamma activity.

The fast gamma oscillations that emerge during sustained photo-activation of SST+ interneurons could reflect the intrinsic synchronisation of SST+ networks, but there are a number of possible scenarios in which this stimulation paradigm could lead to the activation of other hippocampal microcircuits involving network excitation. Depolarising GABA could contribute to recruitment of postsynaptic targets, but perforated patch recordings from hippocampal cells in stratum pyramidale (aCSF only) showed that they were hyperpolarized by light illumination (Supplementary Fig. 5f-g). Alternatively, network excitation and oscillogenesis could emerge following depolarisation block of SST+ interneurons, and subsequent disinhibition, but direct photo-inhibition of SST+ interneurons was not able to generate *de novo* oscillations (Supplementary Fig. 5a-b). However, the power of the light-induced oscillations was markedly reduced following block of either fast excitation or inhibition, and whole cell recordings in putative principal cells indicated that they received weak excitatory postsynaptic currents (EPSCs) throughout light illumination (Supplementary Fig. 5c-e, h-j). This suggests that the light-induced oscillations recorded in LFP do not emerge solely from the activity of SST+ interneurons.

To directly test if photo-excitation of SST+ interneurons leads to a dominant effect of depolarisation block during ongoing gamma oscillations, and whether photo-excitation is associated with net increases or decreases in the spiking activity of other neurons in the network, we performed MEA recordings. We found that SST+ interneurons (spike width: 0.69 +/- 0.03 ms) displayed increased activity throughout the course of step stimulation (Median sustained activation index [IQR] = 0.90 [0.76, 0.98], Z=2346, p<0.001, n=68, one-sample Wilcoxon signed rank test; Fig. 6j-top), and faithfully followed the 8 Hz sine stimulation (Median rank correlation coefficient [IQR] = 0.63 [0.52, 0.72], Z=1484, P<0.001, n=54, one-sample Wilcoxon signed rank test; Fig. 6j-middle). All but 3 of the SST+ interneurons recorded were significantly phase-coupled to the induced fast gamma-frequency oscillations (p<0.05, Rayleigh test), with a mean spike phase of −2.0 [−2.1, −1.8] radians (second-order mean [95% confidence intervals]; n=65; Fig. 6j-bottom). The RS and FS cells showed significantly weaker modulation (Fig. 6j), but did not appear to be suppressed as in the PV-ChR2 experiments, and rather showed an insignificant trends towards both increased activity during step illumination (Median sustained activation index [IQR]; RS: 0.12 [−0.08, 0.54], Z=50, p=0.13, n=11; FS: 0.49 [−0.14, 0.59], Z=24, p=0.09, n=7; one-sample Wilcoxon signed rank tests) and positive correlations with theta-frequency changes in light intensity (Median rank correlation coefficient [IQR]; RS: 0.26 [−0.07, 0.53], Z=49, p=0.16, n=11; FS: 0.28 [0.09, 0.57] (Z=25, p=0.06, n=7, one-sample Wilcoxon signed rank tests). The majority of RS (8/11) and FS cells (4/7) were also significantly phase-coupled to the light-induced fast gamma oscillations (p<0.05, Rayleigh test), but did not show a consistent mean firing phase (RS: F(2,8)=2.3, p=0.18; FS: F(2,2)=4.5, p=0.19; parametric second-order analysis (Zar, 1999)). Overall, this suggests that the dominant change in the network during the induction of fast gamma oscillations is a robust increase in the spiking of SST+ interneurons.

To explore whether the recruitment of SST+ interneurons might differ between step and theta stimulation, we analysed the maximum spike rates in the second half of the stimulation trials (20 ms bins). The maximum spike rates during theta stimulation were significantly higher than during the step stimulation (Z=148, p<0.001, n=54; Wilcoxon signed rank test). As theta stimulation induced faster gamma oscillations than step stimulation (see Fig. 6e), this further suggests that the frequency of fast gamma oscillations depends on the overall levels of SST+ interneuron excitation.

## Discussion

Gamma oscillations depend on synchronised synaptic inhibition, and there is a wealth of evidence suggesting that perisomatic-targeting PV+ interneurons provide the critical inhibitory output for both current and rhythm generation (Mann *et al*., 2005; Bartos, Vida & Jonas, 2007; Oren, Hájos & Paulsen, 2010; Tukker *et al*., 2013; Cardin, 2016; Sohal, 2016; Penttonen *et al*., 1998b). Here, we used optogenetic manipulation of PV+ and SST+ interneurons to explore whether PV+ interneurons have a selective role in gamma rhythmogenesis in the hippocampal CA3 *ex vivo*. Our findings suggest that optogenetically disrupting interneuronal activity, via either photo-inhibition or photo-excitation, generally leads to a decrease in the power and increase in the frequency of ongoing cholinergically-induced slow gamma oscillations. This suggests that both PV+ and SST+ interneurons play key roles in maintaining slow gamma oscillations, and the key differences were that (i) cholinergically-induced gamma oscillations were more readily disrupted by photo-inhibition of SST+ interneurons rather than PV+ interneurons, (ii) manipulation of SST+ interneurons modulated gamma oscillation frequency more robustly than that of PV+ interneurons, and (iii) photo-stimulation of SST+ interneurons could also induce *de novo* fast gamma oscillations.

Slow gamma oscillations in the hippocampal CA3 appear to be generated by synaptic feedback loops between excitatory pyramidal neurons and perisomatic-targeting interneurons, both in brain slices (Fisahn *et al*., 1998; Hajos, 2004; Mann *et al*., 2005; Oren *et al*., 2006; Butler, Hay & Paulsen, 2018) and *in vivo* (Bragin *et al*., 1995; Csicsvari *et al*., 2003; Fuchs *et al*., 2007). In such feedback loops, the period of the oscillation largely reflects the effective time course of inhibitory postsynaptic potentials in the pyramidal cells, which should become shorter with smaller compound inhibitory synaptic currents and/or increased pyramidal cell excitability. The amplitude of the oscillation recorded in the LFP also reflects the amplitude of phasic inhibitory currents in pyramidal neurons (Mann *et al*., 2005; Oren, Hájos & Paulsen, 2010), and during spontaneous gamma oscillations there is a strong correlation between the instantaneous period and amplitude of each gamma cycle (Atallah & Scanziani, 2009). One might thus expect disinhibition to decrease the amplitude and increase the frequency of gamma oscillations, which is largely what we observed with photo-inhibition of either PV+ or SST+ interneurons. Observing similar effects with photo-stimulation of interneurons might be somewhat more surprising. However, our interpretation is that this also effectively disrupts synchronisation within synaptic feedback loops, by silencing a subpopulation of pyramidal cells and thus ‘knocking out’ part of the gamma oscillating network. In both types of optogenetic manipulation, we could thus be recording the activity in residual parts of the network that can maintain synaptic feedback loops, albeit with weaker synchronised inhibition.

While photoinhibition of either PV+ or SST+ interneurons was able to disrupt cholinergically-induced gamma oscillations, it was necessary to use high-powered laser illumination of PV+ interneurons to consistently reduce gamma power, and the oscillations were not abolished under our stimulation paradigms. This is not inconsistent with PV+ interneurons playing a key role in the synaptic feedback loops generating gamma oscillations in the hippocampal CA3, as such a microcircuit should resist disinhibition. Indeed, strong laser illumination was necessary to biochemically silence PV+ interneuron terminals (El-Gaby *et al*., 2016), and thus break this feedback loop. The lack of consistent effects on the frequency of cholinergically-induced gamma oscillations may also be due to the difficulty in silencing PV+ interneurons. Alternatively, it is possible that the remaining PV+ interneurons take longer to fire in each gamma cycle due to the Arch-induced hyperpolarisation. Combined with a more excitable pyramidal neurons, which recover more rapidly form synaptic inhibition, this could leave the overall oscillation frequency unchanged.

It was recently suggested that SST+ interneurons, but not PV+ interneurons, contribute to the generation of slow gamma oscillations in V1 (Chen *et al*., 2017; Veit *et al*., 2017; Hakim, Shamardani & Adesnik, 2018). Our results do not support an exclusive role for SST+ interneurons in slow hippocampal gamma oscillations, but are consistent with an important role for SST+ interneurons in gamma rhythmogenesis across cortical circuits. However, SST+ interneurons largely target the dendritic domains of pyramidal cells, and thus it remains difficult to see how they could directly contribute to the precise timing of pyramidal cell spiking during fast brain oscillations, such as cholinergically-induced gamma oscillations in hippocampal CA3. SST+ bistratified interneurons have been proposed to have similar properties to fast spiking PV+ interneurons, and also form a small portion of synapses close to the soma (Somogyi & Klausberger, 2005; Muller & Remy, 2014), but have been reported to exhibit decreased GABA release under cholinergic stimulation (Gulyás *et al*., 2010). It therefore seems likely that the importance of SST+ interneurons to the generation of cholinergically-induced hippocampal gamma oscillations lies in their modulation of perisomatic feedback loops, via effects on both pyramidal neuron excitability, and the spike rate and precision in PV+ interneurons (Savanthrapadian *et al*., 2014).

While optogenetic manipulation of SST+ interneurons consistently disrupted slow gamma oscillations, we found that photo-stimulation of SST+ interneurons could also induce *de novo* fast gamma oscillations. These GABAergic interneurons should provide a powerful source of circuit inhibition (Somogyi & Klausberger, 2005; Pfeffer *et al*., 2013; Taniguchi *et al*., 2011; Leão *et al*., 2012; Lovett-Barron *et al*., 2012, 2014; Royer *et al*., 2012; Urban-Ciecko & Barth, 2016), but we found that sustained photo-stimulation of SST+ interneurons did not significantly inhibit the activity of STT-neurons, and that pulsed stimulation could drive network excitation. It may be that the hyperactivation of a dense plexus of SST+ processes in the dendritic layers leads to bystander effects on nearby SST-neurons, possibly via ephaptic interactions and/or changes in the extracellular ionic environment (Anastassiou *et al*., 2011; Ferenczi *et al*., 2016b), which counteracts the effects of synaptic inhibition. The generation of fast gamma oscillations appeared to depend on the maintenance of network excitability, as the oscillations were attenuated by block of iGluR. However, the spiking of the majority of STT-RS neurons was only weakly coupled to the phase of light-induced fast gamma oscillations, and without a consistent population spike phase preference, while light-sensitive putative SST+ interneurons showed reliable phase-locking. This could be consistent with fast gamma oscillations representing rhythmic dendritic inhibition from STT+ interneurons, with only weak effects on the spike rate and timing of other neurons in the network.

The mechanism by which a network of SST+ interneurons might generate fast gamma oscillations remains obscure. In neocortex, SST+ interneurons avoid inhibiting each other (Pfeffer *et al*., 2013), although there is evidence for sparse synaptic interactions between SST+ interneurons in the hippocampus (Savanthrapadian *et al*., 2014), and for more generic coupling via gap junctions (Baude *et al*., 2007). More experiments are required to resolve the mechanisms by which optogenetic manipulation of interneurons influences hippocampal gamma oscillations, and whether STT+ neurons contribute to fast hippocampal gamma oscillations during theta and non-theta states *in vivo* (Sullivan *et al*., 2011). However, our findings suggest that SST+ interneurons exert powerful control over the power and frequency of slow hippocampal gamma oscillations, and can switch the network between slow and fast gamma states.

## Materials and Methods

### Transgenic mice

All procedures were performed according to the United Kingdom Animals Scientific Procedures Act (ASPA) 1986 and the University of Oxford guidelines. Adult (older than 8 weeks, both male and female) PV-cre (B6;129P2-Pvalbtm1(cre)Arbr/J), PV-cre-Ai9 (PV-Cre x Gt ROSA (CAG-tdTomato) Hze/J), and SST-cre mice (Sst tm2.1(cre)Zjh/J) were used for all experiments.

### Stereotaxic viral injections

Anaesthesia was induced in mice with 4% isoflurane/medical oxygen mixture (2 L per min). The area around the head was shaved and cleaned in preparation for scalp incision. Anaesthesia was subsequently maintained using 1.5 - 2.5% isoflurane at a rate of 2 L per min. Before the onset of the procedure a cocktail of systemic peri-operative analgesics (Metacam 1 mg/Kg and Vetergesic 0.1 mg/Kg) and a local analgesic (Marcaine 10mg/Kg) were administered subcutaneously (Oxford University Veterinary Services). Following, antibiotic solution was applied on the head and an incision of the scalp was performed that allowed a small craniotomy to be made. A 33/34-gauge needle was attached on a Hamilton Microliter Syringe and used to inject the virus solution at a rate of ≈100 nL/min (viral concentration ≈ 10^12^ genome copies per mL). After every injection, the needle was left stationary for at least three minutes to allow diffusion of virus in the surrounding area. The virus solution was injected with the aid of a stereotaxic frame into ventral CA3 area of hippocampus (2.7 mm caudal and 2.75 mm lateral from Bregma). A total of 600 - 800 nL were injected at two depths (300 - 400 nL at 3.1 mm and 300 - 400 nL at 2.7 mm). Following the injection, local analgesic (Marcaine 10 mg/Kg) was applied on the incised scalp before it was sutured. The animals were then transferred in a heating chamber and allowed to recover. The animals were monitored, and welfare scored in the following days to ensure that they properly recovered after surgery. Injected mice were assessed for viral expression after a minimum of 3 weeks. All viral constructs were acquired from Vector Core Facilities, Gene Therapy Centre (North Carolina, UNC). Viral constructs used: AAV5-EF1a-DIO-ChR2-mCherry, AAV5-EF1a-DIO-ChR2-eYFP, AAV-EF1a-DIO-Arch3.0-EYFP, AAV-Ef1a-DIO-hChR2(E123T-T159C)-p2A-mCherry-WPRE (Dr. Karl Deisseroth), and AAV-CAG-FLEX-ArchT-GFP (Dr. Ed Boyden).

### *Ex vivo* brain slice preparation

Mice were anaesthetised using 4% isoflurane (Oxford University Veterinary Services) and were sacrificed by decapitation after the pedal reflex was abolished. Brains were extracted in warm (30 - 35 °C) sucrose solution (34.5 mM NaCl, 3 mM KCl, 7.4 mM MgSO_4_.7H2O, 150 mM sucrose, 1 mM CaCl_2_, 1.25 mM NaH_2_PO_4_, 25 mM NaHCO_3_ and 15 mM glucose) and transverse hippocampal slices of 350 μm thickness were cut using a Leica vibratome (VT 1200S) (Huang et al. 2013). Slices were then immediately placed in an interface storing chamber containing warm (30 - 35 °C) aCSF (126 mM NaCl, 3.5 mM KCl, 2 mM MgSO_4_-7H_2_O, 1.25 mM NaH_2_PO_4_, 24 mM NaHCO_3_, 2 mM CaCl_2_ and 10 mM glucose) at least one hour to equilibrate. All solutions were bubbled with 95% O_2_ and 5% CO_2_ beginning 30 minutes before the procedure until the end of the experiment.

### Electrophysiology

Extracellular recordings were conducted in an interface recording chamber at 33-34 oC. Visualisation of the slices and electrode placement was performed using a Wild Heerbrugg dissection microscope. Local field potentials were recorded by inserting a borosilicate glass electrode filled with aCSF (tip resistance = 1 - 5 MΩ) in CA3 pyramidal layer. Data were acquired and amplified (x 10) by Axoclamp 2A (Molecular Devices). The signal was further amplified x 100 and low pass filtered at 1 KHz (LPBF-48DG, NPI Electronic). The signal was then digitised at 5000 samples per second by a data acquisition board (ITC-16, InstruTECH) and recorded from the IgorPro (Wavemetrics). Gamma oscillations were induced by the application of 5 μM carbachol (Cch). The LFP signal was quantified using real-time fast Fourier transform (FFT) analysis and oscillations were detected by a peak in the power spectrum at low-band frequencies (25 Hz - 49 Hz). For unit recordings a linear 16 channel tungsten multi-electrode array (MEA; MicroProbes) was lowered in the CA3 subfield. The array channels had 100 μm spacing to ensure full coverage of the hippocampus. The MEA was mounted on an RHD2132 Amplifier board and connected to the RHD2000 USB Interface Board (Intaan Technologies). Data were acquired at a rate of 20000 samples per second using the RHD2000 rhythm software (Intaan Technologies).

Intracellular recordings were always conducted in a single submerged chamber (26 - 32 oC) using borosilicate glass pipettes (5-12 M). The signal was acquired through the MultiClamp 700B amplifier (Molecular Devices) and digitised at a rate of 10000 samples per second by a data acquisition board (ITC-18, InstruTECH) and was then recorded using the Igor Pro software. The signals were low pass filtered (Bessel) at 10 kHz for current clamp mode and 3 kHz for voltage clamp (VC) mode. Slice and cell visualisation were achieved using oblique illumination and monitored through a HAMATATSU ORCA - ER digital camera. Filtered white LED (460 +/- 30 nm, 1.53 mW, Thor Labs) via epi-illumination was used to activate channelrhodopsin (ChR2). Filtered white LED (525 +/-20 nm, 1.45 mW, Thor Labs) via epi-illumination was used to activate archaerhodopsin (Arch). For a power of 1.5 mW, the illuminated area is 3.68 mW / mm^2^. Cell attached recordings were performed in current clamp (IC) mode (Multiclamp software) using glass pipettes filled with aCSF. For whole cell current clamp recordings pipettes were filled with internal solution containing 110 mM KGluconate, 40 mM HEPES, 2 mM ATP-Mg, 0.3 mM GTP-NaCl, 4 mM NaCl, (3-4 mg/ml biocytin, Sigma). For whole cell voltage recordings pipettes were filled with internal solution containing 140 mM Cesium methanesulfonate, 5mM NaCl, 10 mM HEPES, 0.2 mM EGTA, 2 mM ATP-Mg, 3 mM GTP-Na, 5 mM QX-314, (3-4 mg/ml biocytin). Series resistance compensation was not performed in all cells included for analysis. For perforated patch recordings the tip of the pipette was filled with a KCl-containing solution (150 mM KCl and 10mM HEPES, pH 7.2-7.3; Osmolality 300 mOsmol/Kg). The rest of the pipette was filled with the same KCl solution containing 5 μM gramicidin D (1:1000 DMSO dilution, Sigma) and 10 μM Fluorescein (Sigma) to visualise if there was spontaneous rupture of the membrane during patching experiments.

### Light delivery

For photo-activation (ChR2) experiments, light illumination was delivered through a fibre optic using a blue LED (470 +/- 20 nm, Thorlabs, M470F3; max power at fibre optic tip = 10 mW). For photo-inhibition (Arch) experiments light illumination was delivered through a fibre optic by a green LED (530 +/- 30 nm, Thorlabs, M530F2; maximum power at fibre optic tip = 4.25 mW) and with an amber LED (595 +/- 20nm, Doric, maximum power at fibre optic tip = 5 mW). LED module output was controlled using the Igor Pro software. Laser photo-inhibition experiments were also performed with a green laser (MatchBox series, 532 +/- 0.5 nm, maximum power at fibre optic that was used approx. 40 mW). In these experiments the data were acquired at a rate of 10000 samples per second using IgorPro. The laser was operated manually, and the light duration was recorded using an Arduino Uno board that created a digital time stamp. Experiments were only included if the laser illumination duration was between 19.6 - 20.7 seconds. The area of light illumination was estimated to have a diameter of 1 - 2 mm and therefore for a power of 10 mW the light intensity was between 0.8 - 3.2 mW / mm^2^.

### Histology and imaging

After electrophysiological recordings, acute brain slices were fixed in 4% PFA overnight. Slices were kept in PBS (Phosphate Buffered Saline: 1.37 mM NaCl, 2.7 mM KCl, 10 mM Na_2_HPO_4_, 2 mM KH_2_PO_4_) at 4 °C for short-term storage. For biocytin labelling the slices were washed with 1X PBS 3-4 times and permeabilized with freshly prepared 0.3%-Triton 1X PBS for 4 - 5 hours. Streptavidin conjugated to Alexa FluorTM 488 (Invitrogen S32355) in PBS-T 0.3% (1:500) was incubated overnight at 4 oC. The slices were then washed 4 - 5 times in PBS for 2 hours. Slices were mounted on glass slides using mounting media (DAKO). Confocal images (1024×1024) were acquired on a Zeiss LSM700 upright confocal microscope using the 10x air objective and digitally captured using the default LSM acquisition software. Pyramidal cell reconstruction was performed on neuron studio and simple neurite tracer plugin on Fiji.

### Analysis of local field potentials

In order to characterise and analyse the oscillations, a hanning window was applied and the power spectra were calculated as the normalised magnitude square of the FFT (Igor Pro). The 50 Hz and 100 Hz frequencies were not included in the analysis to exclude the mains noise and its harmonic component. The oscillation amplitude was quantified firstly by measuring the peak of the power spectrum termed as peak power and secondly by measuring the area below the power spectrum plot in the gamma-band range (20 - 100 Hz) termed as power-area. The peak frequency of the oscillation was obtained by measuring the frequency at which the peak of the power spectrum occurred in the gamma-band range. In order to quantify when Cch-induced oscillations where abolished upon light stimulation and to exclude the peak frequencies of those oscillations from further analysis one of the two criteria had to be met. Firstly, an auto-correlation of the oscillations was computed and was fitted with a Gabor function (*f*(*x*) = (*A* * *cos* (2*π* * *f* * *x*)) * *e* – *x*2/2 * *tau*). The first criterion was met if the resulting Gabor fit had a linear correlation coefficient, r > 0.7 and product of 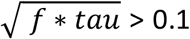 (> 0.15 for frequencies higher than 50 Hz). The second criterion was a power-area bigger than 125 μV^2^ in the range of +/- 5 Hz of the peak frequency. The power-area was always included in the analysis even if oscillations were abolished. The power spectrum analysis for *de novo* oscillations was performed in the range of 52 – 149 Hz with the only criterion for oscillation presence being that the power-area in +/- 5 Hz of the peak frequency was larger than 40 μV^2^. Hilbert transforms were used to obtain instantaneous gamma magnitude for sinusoidal modulation of gamma oscillations (band-pass filtered 20 - 120 Hz). For visualisation purposes the magnitude of the continuous wavelet transform was used normalised by max value (Morlet wavelet; nondimensional frequency = 6).

### Spike detection and analysis

Unit detection was performed using custom-written procedures in MATLAB (Mathworks). Extracellular spikes from the 16 channel MEA were detected as described before by Quiroga and colleagues (Quiroga, Nadasdy & Ben-Shaul, 2004; Quian Quiroga, 2009). Briefly, the MEA data were processed with an elliptical band-pass filter (for spike detection: 4th order, 300 - 3000 Hz, for spike sorting: 2nd order, 300 - 6000 Hz). Spikes were detected as signals exceeding 5 standard deviation (s.d.) of the noise 5 ∗ *σn* = *median* {|*x*| / 0.6745}. Signals that exceeded 10 times the s.d. of the detected spike amplitudes were eliminated as artefacts/population spikes. Subsequently, spikes that had peaks occurring at the same time (< 0.1 ms) across channels were grouped together as one unit. This prevented detection of the same unit more than once. Clustering of the detected spikes was performed using custom-written procedures in Igor Pro. A spike sorting procedure adapted from Fee and colleagues was used to explore whether neurons displaying specific spike waveforms were selectively recruited by optogenetic stimulation (Fee, Mitra & Kleinfeld, 1996). Briefly, spike metrics were converted into z scores, over-clustered using an in-built k-means algorithm, and progressively aggregated if the intercluster distance was <2.5 and merging did not produce more violations of refractory period of 2 ms. Analysis was performed on the clustered spikes, with auto-correlation and cross-correlation plots used to validate the clustering procedure. Spike metrics from the average waveform for each cluster were used to identify different waveform types via a k-means algorithm. This clustering procedure is likely to be conservative, and underestimate the firing rate of individual neurons, but was deemed sufficiently robust to detect any bias in optogenetic recruitment. A single unit cluster was identified if it i) had less than 1.4% of its total spike waveforms within 2 ms of its refractory period and ii) consisted of more than 800 members. When a cluster did not obey these criteria, it was merged with other clusters that had similar action potential waveforms giving rise to a multi-unit cluster.

Clusters were identified as expressing ChR2 if the spike rate in the first 100 ms of the step stimulus was 3 s.d. above the baseline spike rate. The remaining clusters were classified based on the the delay between the negative and positive peaks in the average waveform as fast-spiking (<0.6 ms) or regular-spiking (>=0.6 ms). The Activation Index was calculated over the last second of the step stimulus as the difference between the light-induced and baseline spikes rates divided by their sum, and designed to measure sustained firing. The Theta Modulation Index was calculated as the rank correlation coefficient between the spike time histogram and the theta-modulated amplitude of the light stimulus.

### Statistics

Repeated measures ANOVA (rmANOVA) was performed in SPSS with Greenhouse-Geisser correction where required (i.e. significance in Mauchly’s test for sphericity) and followed by Bonferroni-corrected post-hoc paired t-tests. Linear correlations, circular correlations, and Bonferroni-corrected one sample t-tests were performed using Igor Pro. Scatter-bar charts were generated using PRISM. Circular statistics of spike phase relative to ongoing oscillations in the LFP were calculated using in-built functions in Igor Pro. The measurements spiking rates deviated from normality, and were analysed using non-parametric statistical tests performed in SPSS: differences between cell types were analysed using Kruskal-Wallis Test, followed by posthoc Dunn’s test, with Bonferroni correction for multiple comparisons. Differences across stimulus types (step and theta) were analysed using the Wilcoxon signed rank test, and the significance of modulation indices analysed using the one-sample Wilcoxon signed rank test (H_0_=0). In all figures, the bar charts display the average value and the error bars represent the standard error of the mean, unless explicitly stated otherwise. Stars represent significance values where * p <0.05, ** p <0.01 and *** p <0.001.

---

**Supplementary Figure 1:**
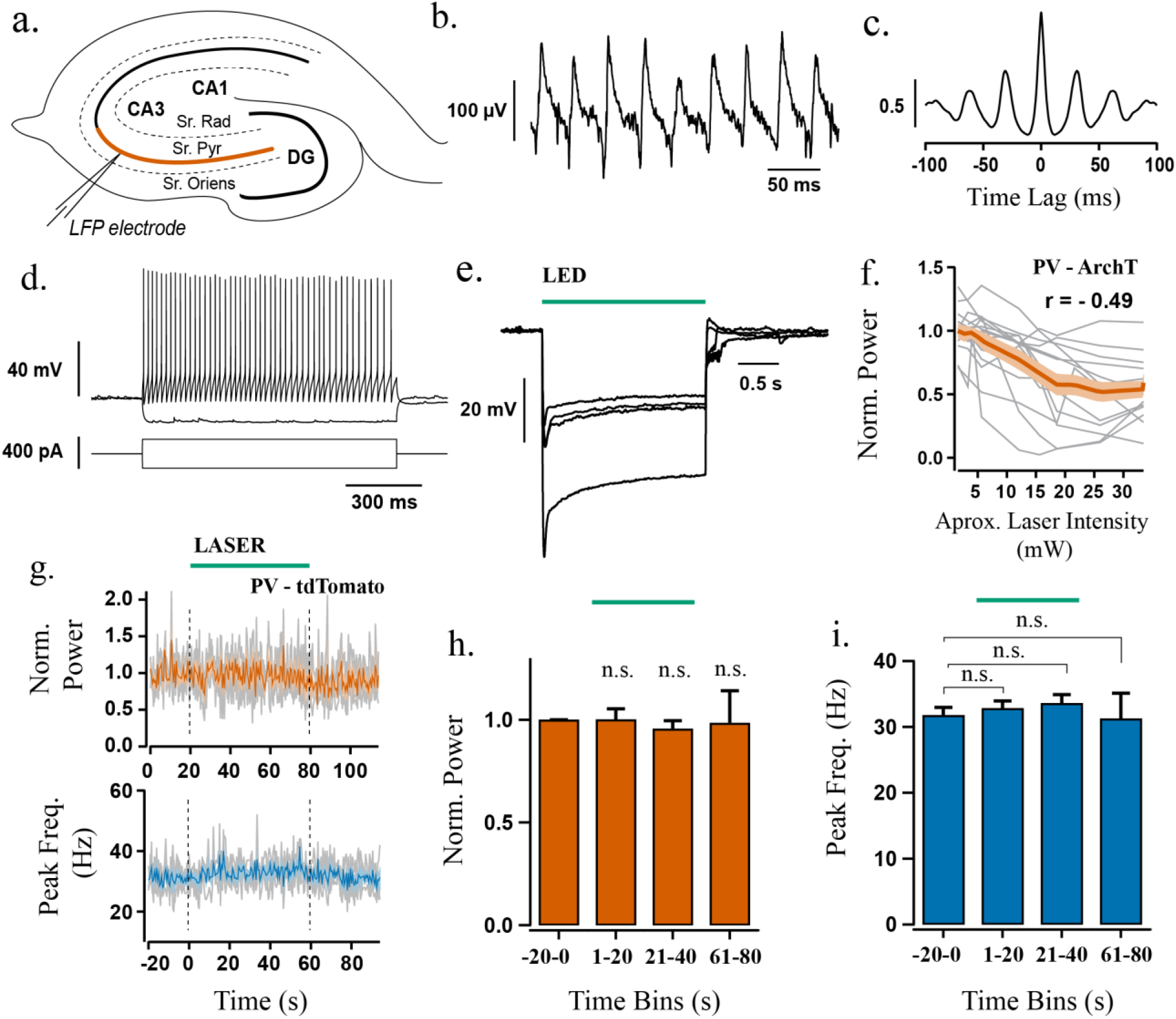
a-c) Recording gamma oscillations in hippocampal CA3. a) Illustration of the electrophysiological setup used to record gamma oscillations using a glass field electrode in stratum pyramidale of hippocampal area CA3 (coloured line indicates the CA3 area where recordings were obtained). b) Representative LFP recording from the ventral CA3 area obtained by the application of 5 μM Cch. c) Autocorrelogram of the recording in b) illustrating that the oscillation is rhythmic with a period of 31 ms. d-e) Validation of functional Arch expression in PV+ neurons. d) Current clamp recording of an ArchT-GFP expressing PV+ cell from CA3 area in response to steps of depolarising and hyperpolarising current injections. e) Potent hyperpolarisation of four PV+ interneurons during green light illumination in aCSF (1.45 mW). f) Power change between the last half of laser stimulation from baseline against approximate light intensity. Grey lines represent single experiments (n = 14). The orange line is the average response and the orange shaded area represents SEM. g-i) Responses to laser illumination in control slices from PV-Ai9 mice. g) Top: Change in power-area normalised to baseline calculated in 1 second bins across experiments (n = 4). Bottom: Peak frequency of the oscillation calculated in 1 second bins across experiments (n = 4). Time between dotted lines indicates the duration of laser illumination (duration of approx. 1 minute and a light intensity of approx. 25-41 mW). h-i) quantification of normalised power and peak frequency. n.s. p >= 0.05. Changes in peak frequency were analysed using rmANOVA, followed by post-hoc paired t-tests with correction for multiple comparisons. Solid brackets represent paired t-tests and standalone star symbols represent one-sample t-test versus normalised baseline. Error bars and shaded area are SEM and coloured line the population mean.

**Supplementary Figure 2:**
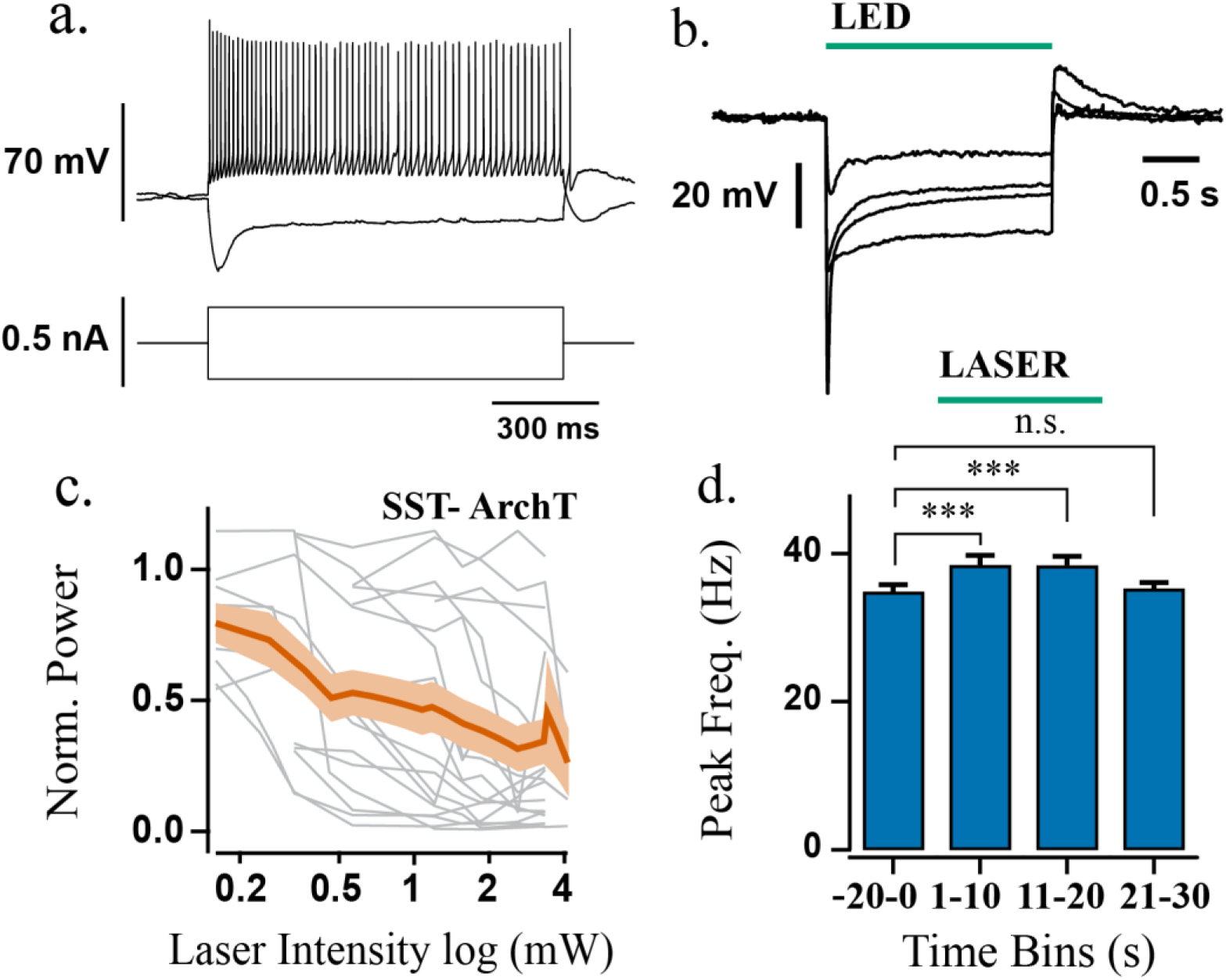
a-b) Validation of functional opsin expression in SST+ interneurons. a) Current clamp recording of an SST+ cell from CA3 area in response to steps of depolarising and hyperpolarising current injections. b) Potent hyperpolarisation of four SST+ interneurons during green light illumination (1.45 mW). c-d) Effects of laser power on network activity. c) Power change between the last half of laser stimulation to baseline against approximate light intensity (n = 17). Coloured line represents the average and shaded area the SEM. d) Average peak frequency for half-maximal response of SST+ interneuron photo-inhibition (light intensity for each experiment that changed power-area by half of the maximum response). rmANOVA, F(3, 48) = 19.27, p < 0.001; ***p < 0.001, rmANOVA followed by post-hoc paired t-test for multiple comparisons. Solid brackets represent paired t-tests.

**Supplementary Figure 3:**
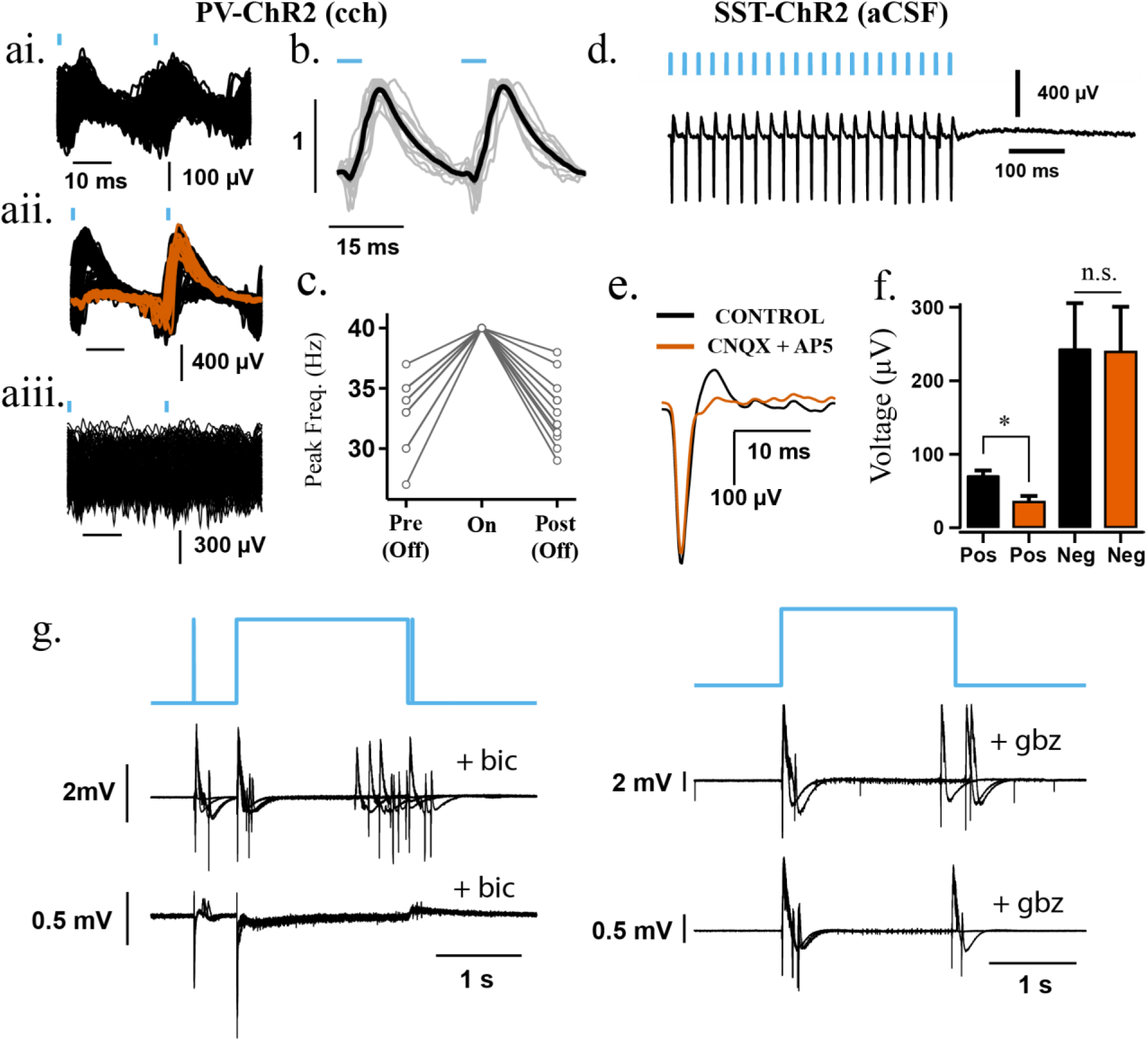
a-c) Responses to pulse stimulation in slices with Cch-induced gamma oscillations from PV-ChR2 mice. a) Extracted waveforms following two consecutive pulses at 40 Hz (1 ms) from one experiment showing i) strong entrainment, ii) entrainment of half of the cycles and inhibition of the other half, and iii) no entrainment. Coloured lines represent a subset of extracted waveforms. b) Average waveform following two consecutive pulses at 40 Hz with pulse widths of 5 ms (n = 13/13). Black line is the population average, grey lines represent individual experiments and dark-grey/error bars shaded area is the SEM. c) Peak frequency of oscillation before (Pre (Off)), during (On) and after (Post (Off)) light stimulation at 40 Hz with pulse widths 5 ms (n = 13/13). Note that the slices with 20 Hz induction were not tested with 5 ms pulse width. d-g) Responses to pulse stimulation in slices from SST-ChR2 mice without cholinergic activation (aCSF). d) Representative LFP recording from CA3 area illustrating field responses during 40 Hz light pulses in the absence of Cch (1ms pulse width-5.5 mW). e) Average waveform obtained with 1 ms pulse width at 40 Hz from one experiment before (black line) and after application of 20 μM CNQX and 40 μM AP5. In one experiment only 20 μM CNQX was applied but exhibited the same effect. f) The magnitude of negative and positive peaks before and after iGluR blocker application (n = 4). iGluR blockers used: 20 μM CNQX, 40 μM AP5, n = 3; 20 μM CNQX, n = 1. Positive peak before (MaxC) and after drug (MaxD) application (t = 3.49, p = 0.03, paired t-test). Negative peak before (MinC) and after drug (MinD) application (t = −0.61, p = 0.58, paired t-test). *p <0.05, n.s. p >= 0.05. g) Induction of epileptiform bursts during photo-stimulation of SST+ interneurons following GABA_A_R blockage (n = 2 at 20 μM bicuculine (bic), n = 2 at 10 μM gabzine (gbz)).

**Supplementary Figure 4.**
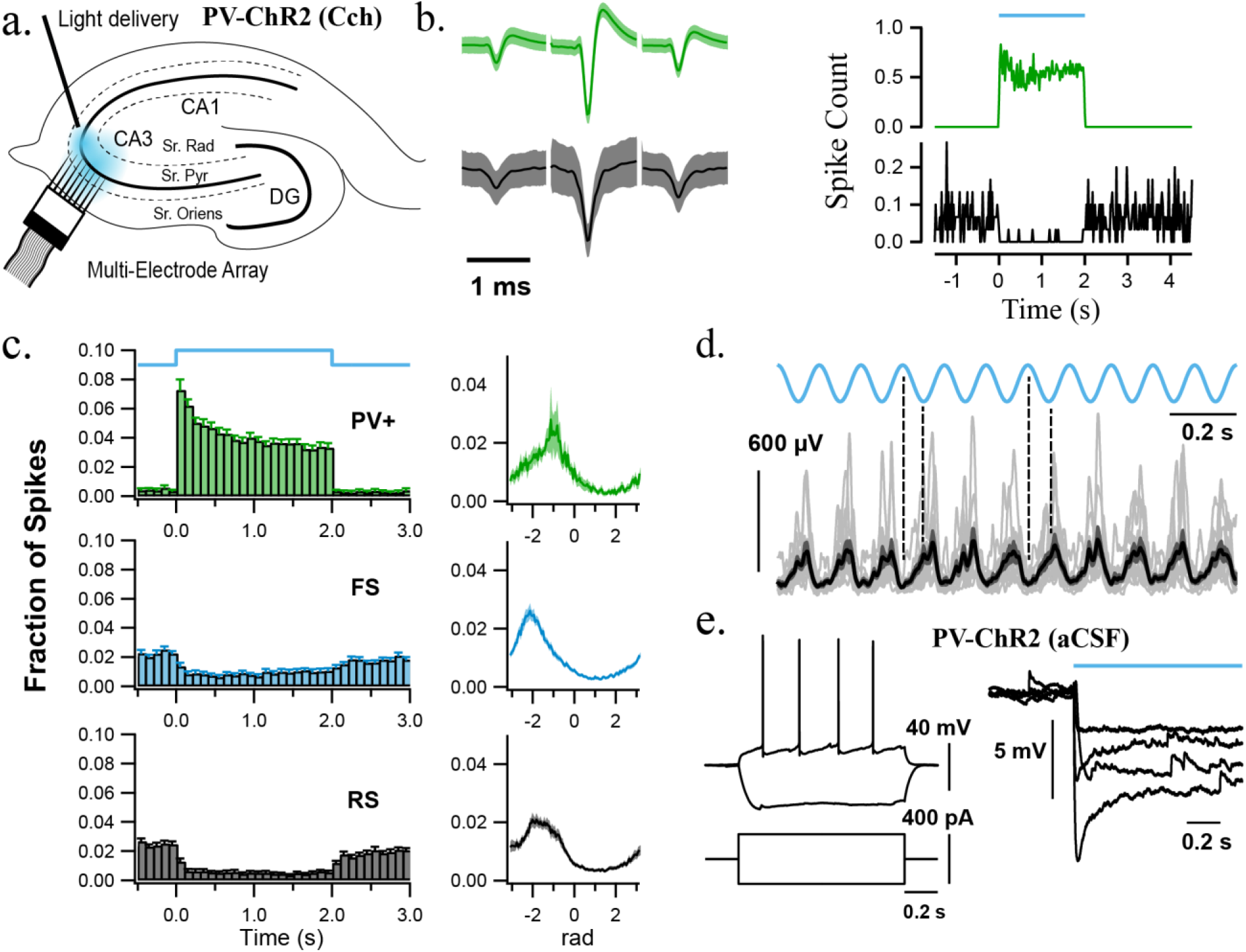
**(a-c):** Multi-unit recordings during PV+ interneuron sustained photo-excitation in hippocampal slices with Cch-induced gamma oscillations. a) Schematic diagram of the hippocampus illustrating MEA recordings during blue light illumination (5.5 mW) in CA3. b) Left - Representative average spike waveforms, Right – spike histograms during sustained light illumination. (green FS single units, black RS multi-unit). c) Mean spike time histograms (left) and spike phase histograms (right) of photo-tagged PV+, FS and RS cells. Shaded regions represent standard deviation. The PV+ interneurons fired at a significantly later phase of the oscillation than the RS cells (F(2,55) = 5.36, p = 0.007, Two-sample Hotelling test). d) Instantaneous amplitude of the Hilbert transform during theta photo-activation (1 mW) overlaid across experiments (grey traces, n = 12), black represents the mean and dark grey the SEM. Dotted lines illustrate that high light intensity decreased gamma power. e) Intracellular recordings from pyramidal cells in aCSF, showing responses to current steps (left) and hyperpolarisation in response to PV+ interneuron photo-activation (n = 4).

**Supplementary Figure 5:**
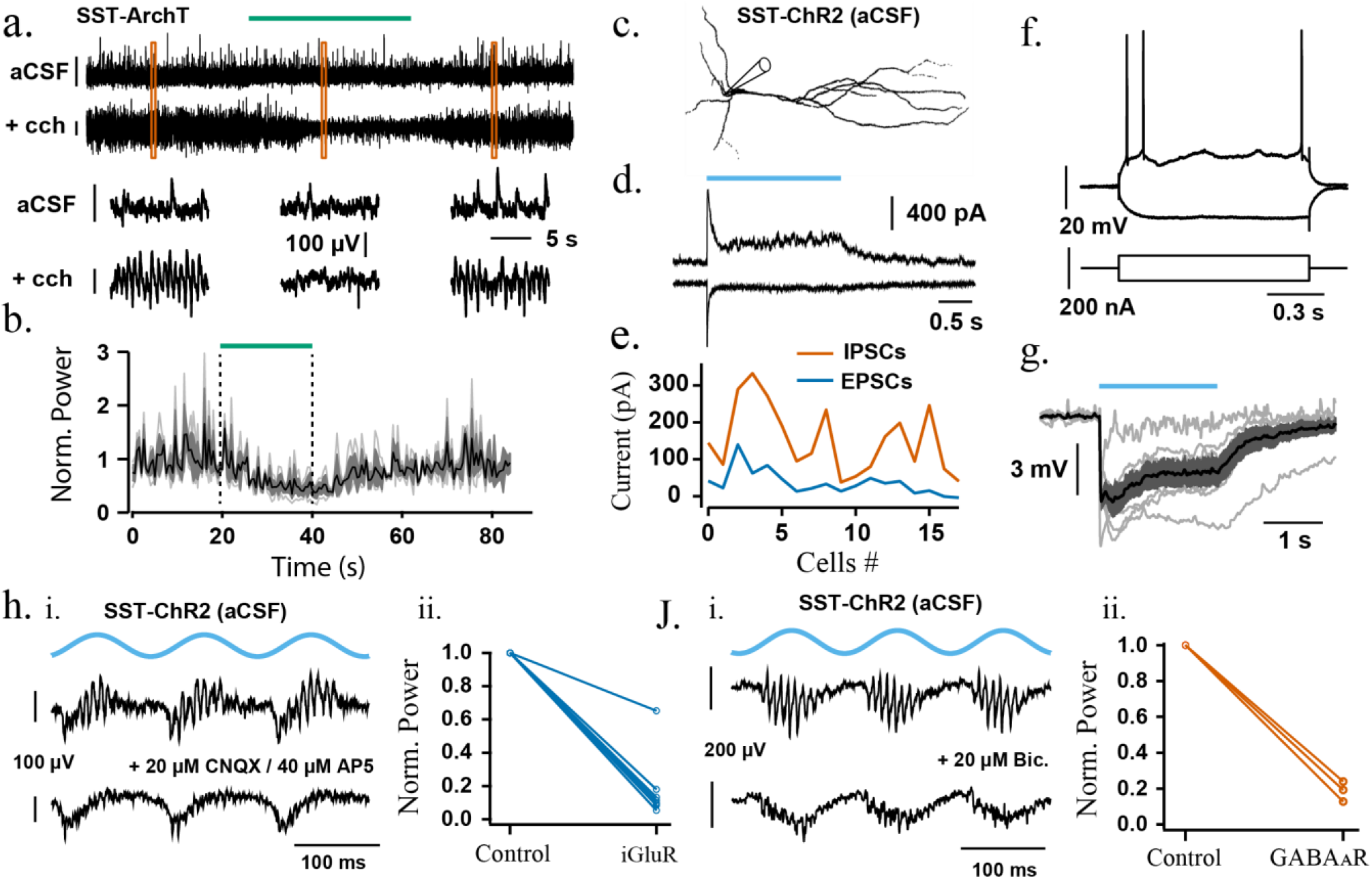
(a–b) Responses to laser illumination in SST-cre mice expressing ArchT-GFP. a) Representative LFP recording with and without the presence of Cch during green laser illumination (approx. 18.6 mW). Orange squares represent the bottom sections of LFP that were magnified. b) Change in power-area normalised to baseline calculated in 1 second bins across experiments (n = 3) in the absence of Cch; Dotted lines indicate the duration of laser illumination. (c-e) Voltage clamp recordings from putative pyramidal cells during photo-activation of SST+ interneurons (1.53 mW). c) Reconstruction of a recorded cell with typical pyramidal cell morphology in CA3 hippocampal area. d) Representative voltage clamp recording of the cell in c) held at 0 mV (top) and −70mV (bottom) to isolate IPSCs and EPSCs, respectively. e) Comparison of EPSCs and IPSCs during SST+ interneuron photo-activation across cells (n = 18). Blue line = EPSCs, Orange line = IPSCs. (f–g) Perforated patch recordings in CA3 pyramidal cell layer in SST-cre mice expressing ChR2-mcherry. f) Current clamp recording in a putative pyramidal cell in response to depolarising and hyperpolarising current injections. g) Hyperpolarisation of the membrane voltage of perforated patched cells (n = 6) during blue light illumination (1.53 mW). Grey traces represent individual cells, black trace the average and dark grey shaded area the SEM. (h-j) pharmacology of de-novo oscillations induced by sinusoidal blue light illumination in SST-cre mice expressing ChR2-mcherry (1 - 10 mW). Representative LFP recording in CA3 before (top) and after (bottom) application of h.i) iGLUR blockers and j.i) GABA_A_R blockers. Power-area change before (control) and after application h.ii) iGLUR blockers and j.ii) GABA_A_R blockers. GluR blockers used: 20 μM CNQX, 40 μM AP5, n = 3; 10 μM CNQX, 20 μM AP5, n = 1; 20 μM CNQX, n = 1; 3mM kynurenic, n = 3. GABA_A_R blockers: 20 μM Bicuculline, n = 2; 20 μM Gabazine, n = 1.

